# Metabolic modeling links leaf anatomy to environment-specific benefits on the C3-C4 spectrum

**DOI:** 10.64898/2026.04.28.721324

**Authors:** Alejandro León-Ramírez, Tiago M. Machado, Sezin Dogan, Andreas P.M. Weber, Urte Schlüter, Nadine Töpfer

**Author notes:** Both authors contributed equally (Alejandro León-Ramírez, Tiago M. Machado).

## Abstract

C4 photosynthesis evolved from the ancestral C3 pathway through coordinated leaf anatomical and metabolic reorganization that concentrates CO_2_ to reduce photorespiration. Quantitative understanding of these structure-function relationships remains limited. Here we used anatomy-aware metabolic modeling of a mesophyll-bundle sheath cell system to analyze the interdependence between leaf anatomy and photosynthetic metabolism on the C3-C4 spectrum. Our model faithfully recapitulates the transitory steps from C3 to C4 photosynthesis, reveals a crucial role for plasmodesmata in enabling the C3 to C4 transition, and points at potential pre-C2 metabolic states that provide benefits under conditions that promote elevated photorespiration. Incorporating bundle cell suberisation with our model predicts reduction of PSII activity and dominance of the NADP-ME C4 subtype in leaves with suberized bundle sheath cells and proposes a role for oxygen evolution at PSII as a potential driver for this mechanism. Varying bundle sheath leakage and photorespiratory conditions along the C3-C4 spectrum identify conditions under which C3-C4 intermediacy provides energetic benefits and underlines the notion of intermediate photosynthesis as a stable evolutionary state. Overall, our study sheds new light on the quantitative relationship between leaf anatomy and metabolism and its interaction with the environment and suggests targets for climate-adaptation in C3 plants.

## Introduction

Increasing temperatures and weather extremes represent major threats to global crop productivity. To sustain future yields, it is essential to develop crop varieties capable of maintaining high photosynthetic efficiency under these anticipated conditions. A promising strategy to achieve this goal, is the introduction of either C3-C4 intermediacy or C4 photosynthesis into C3 crops. Evolution has shaped these specialized pathways to optimize carbon fixation efficiency and mitigate the energetic penalties associated with photorespiration, particularly under warm conditions. C4 photosynthesis relies on the spatial separation of initial CO_2_-fixation and subsequent CO_2_ reduction between mesophyll and bundle sheath cells, thereby concentrating CO_2_ around rubisco. This specialization is underpinned by extensive anatomical and metabolic modifications. C4 plants exhibit Kranz anatomy, in which mesophyll cells form a concentric layer around enlarged bundle sheath cells. Tight cell-to-cell contact and high plasmodesmata density between these cell types facilitate efficient metabolite exchange (Danila et al. 2016, Danila et al. 2018). Although the C4 pathway incurs higher ATP costs than C3 photosynthesis, these expenditures are offset by a significant reduction in photorespiratory flux (Langdale 2011; Yin and Struik 2021). The magnitude of this trade-off varies among C4 subtypes—distinguished by their primary bundle sheath decarboxylation enzymes (NADP-ME, NAD-ME, or PEPCK)—and is further influenced by structural features such as the degree of suberisation of bundle sheath cell walls (Langdale 2011; Yin and Struik 2021). C3 or C4 photosynthesis is not a binary syndrome but operates on a spectrum. C3-C4 intermediates of type I, also referred to as type I C2 plants, reduce photorespiration through a glycine shuttle that recycles photorespiratory metabolites and concentrates carbon around bundle sheath rubisco, thereby improving photosynthetic efficiency and carbon assimilation in warmer and dryer environments. Some species additionally operate a partial C4 cycle and are classified as type II C2 plants.

Comparative studies indicate that anatomy drives the evolution of C4 biochemistry and not the reverse. C4 photosynthesis arose repeatedly in lineages whose C3 members were already preconditioned by enlarged bundle sheath cells and closely spaced veins, which favour the refixation of photorespired CO_2_ (Sage 2004, Christin et al. 2013, Griffiths et al. 2013). Implementing C3-C4 or C4-like traits in C3 plants therefore requires reorganization of leaf anatomy, intercellular coordination, and metabolic regulation (Leegood 2002, Schuler et al. 2016). Prior to undertaking such bioengineering efforts, it is essential to elucidate the anatomical, metabolic, and regulatory determinants that are both necessary and sufficient to support the emergence of an efficient heat- and drought-adapted photosynthetic apparatus.

Computational modeling is increasingly valuable in this context, enabling quantitative assessment of energy fluxes, metabolite partitioning, and carbon assimilation to guide the rational design of C4-like traits in C3 species. For example, the Farquhar–von Caemmerer–Berry model and its extensions provide powerful tools to simulate photosynthetic performance and to evaluate trade-offs between ATP demand and CO_2_ fixation rates (Caemmerer 2000, Yin et al. 2021). While these biochemical models capture enzyme kinetics and gas exchange at the leaf level, they cannot fully represent the underlying metabolic complexity or intercellular coordination of intermediate and C4 photosynthesis. To address this, large-scale stoichiometric modeling approaches have been developed to predict system-wide flux distributions across cell types. Among the first flux balance models of C4 photosynthesis, the C4GEM framework introduced a cell-type specific model, with mesophyll and bundle sheath submodels, in which predefined decarboxylation enzyme activities successfully recapitulated the flux patterns characteristic of the major C4 subtypes (Dal’Molin et al. 2010). Building on this foundation, two-cell-type modelling approaches have since been refined and widely applied as reviewed here (Machado et al. 2025). These models have provided important insights into the evolutionary trajectories underlying the C3-to-C4 transition. Mallmann et al. (2014) coupled a flux-balance model to a biochemical model of leaf photosynthesis (Heckmann et al. 2013; Caemmerer 2000) to investigate the role of photorespiration in driving C4 evolution. Blätke and Bräutigam (2019) studied metabolic fluxes in a two-cell-type model across varying environmental conditions, and identified resource limitation alongside high photorespiratory flux as a key driver of the C3-to-C4 transition. They further investigated how light availability and its distribution between cell types influence the selection of decarboxylation enzymes, thereby implicitly accounting for the anatomical specializations of C4 leaves. Yet, although leaf photosynthetic diversity is governed by a tight coupling between form and function, a systematic analysis of the structure–function relationship between leaf anatomy and metabolic processes remains lacking.

Here we develop an anatomy-aware flux-balance model of an integrated mesophyll–bundle sheath system to investigate the structure–function relationships underlying photosynthetic diversity across the C3-C4 spectrum. By incorporating key anatomical constraints, such as mesophyll-to-bundle sheath volume ratios, plasmodesmatal connectivity, and bundle sheath suberisation, the model enables hypothesis-driven exploration of how anatomical specialisation shapes photosynthetic metabolism. The model recapitulates known features of C3-C4 biochemistry, predicts previously unrecognized energetically favorable pre-C2 states under elevated photorespiration, and supports the hypothesis that minimizing O_2_ evolution in the bundle sheath is a major driver of PSII downregulation and the emergence of NADP-ME dominance following cell wall suberisation.

## Results

### Constructing an anatomy-ware model of mesophyll and bundle sheath cell metabolism

Starting from a core model of plant metabolism (Shameer et al. 2018), we constructed a mesophyll-bundle sheath system by duplicating the metabolic network to represent each cell type and linking those via plasmodesmatal metabolite transport. Gas exchange with the intracellular air space was restricted to the mesophyll and nutrient exchange with the vasculature was limited to bundle sheath cells (Supplementary Information, Model reconstruction, Supplementary Tables S1 - S6). To represent ambient conditions, we constrained rubisco in both cell types to a default carboxylation-to-oxygenation ratio (v_c_:v_o_) of 3:1 (Gutteridge and Pierce 2006; Cheung et al. 2014). Additionally, we modelled the carbon-concentrating mechanism around bundle sheath rubisco by allowing a second bundle sheath rubisco population to operate without oxygenase activity using only CO_2_ provided through decarboxylation reactions, as previously demonstrated (Blätke and Bräutigam 2019). This computational artifice allows the model to balance the cost associated with photorespiration against the cost of running a C4 cycle for carbon concentration in the bundle sheath cells. The resulting two-cell-type model comprises 1995 reactions and 1724 metabolites distributed across 6 organelles and the cytosol in each cell type (Fig. 1).

**Figure 1:**
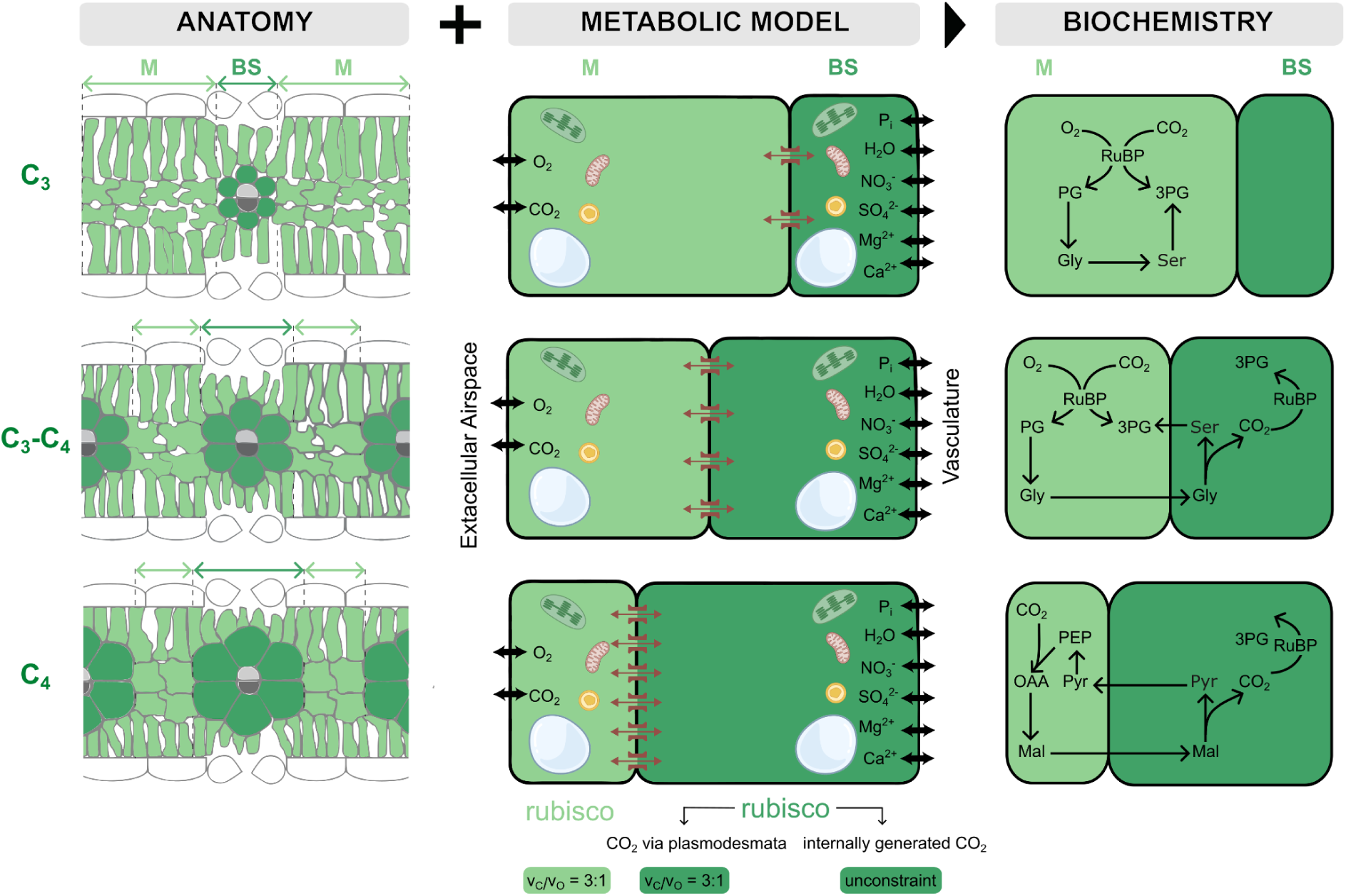
Simulating the structure-function relationship in an integrated model of mesophyll and bundle sheath cell metabolism. Left) Anatomical constraints for the metabolic model were derived by approximating mesophyll and bundle sheath volumes and plasmodesmatal connectivity from leaf sections and anatomical measurements of C3, intermediate, and C4 species. Middle) In the two-cell-type model the mesophyll and the bundle sheath cells are connected via plasmodesmatal transport reactions. The mesophyll cells exchange gases with the extracellular airspace; the bundle sheath cells exchange nutrients with the vasculature. Increasing plasmodesmatal connectivity along the C3-C4 spectrum is modeled through cell-type volume ratio-dependent weighting of transport fluxes. Rubisco carboxylation:oxygenation ratio is initially set to 3:1 in both cell types. A second bundle sheath rubisco population fixes CO_2_ from internal decarboxylation reactions without oxygenase activity. Right) Integration of the anatomical constraints with the metabolic model predicts photosynthetic metabolism on the C3-C4 spectrum. Abbreviations: 3PG - 3-phosphoglycerate, Gly - glycine, MAL - malate, OAA - oxaloacetate, PG - 2-phosphoglycolate, Pyr - pyruvate, RuBP - ribulose 1,5-bisphosphate, Ser - serine.

We accounted for differences in leaf anatomy along the C3-C4 spectrum by imposing two interdependent data-informed anatomical constraints: the bundle sheath to mesophyll volume ratio and the plasmodesmatal connectivity between the two cell types. To define ranges of biologically relevant bundle sheath to mesophyll volume ratios (V_BS/M_) we collected leaf cross-sectional and paradermal imaging data of 84 mono- and dicot species, of which 25 were C3, 28 intermediates, and 31 C4 plants and estimated the leaf volume enriched in mesophyll and bundle sheath cells (not the volume of individual cells) from the images. This resulted in interquartile ranges for V_BS/M_ of 0.5 - 1.1 for C3, 0.7 - 1.7 for intermediates, and 1.2 - 2.2 for C4 species. All exchange fluxes between mesophyll and bundle sheath cells were scaled according to these ratios. Since the interface available for plasmodesmata as well as the plasmodesmata density increases with V_BS/M_ (Danila et al., 2016) plasmodesmatal fluxes were additionally weighted with a function α(V_BS/M_) (Materials and Methods and Supplementary Information, Anatomical constraints, Supplementary Fig. S1 - S3). Given the interdependencies of these two anatomical specializations we refer to the simultaneous change of these parameters as changes to the BS-M anatomy, where a given value denotes the underlying V_BS/M_-ratio.

For model analysis we assumed agricultural settings and simulated carbon-limiting conditions with a maximal CO_2_ uptake rate of 20 µmol m^-2^ s^-1^ and a light intensity (PPFD) of 1000 µmol photons m^-2^ s^-1^ (Amthor et al. 2019). Photon uptake was modeled independently for each cell type with maximum rates determined by incident PPFD. Cellular maintenance was modelled as described in (Töpfer et al. 2020). We simulated leaf metabolism by maximizing growth subject to minimal enzyme investment which was approximated by total reaction flux. Flux solutions were computed using weighted parsimonious flux balance analysis (wpFBA) (Materials and Methods and SI, Anatomical constraints). This approach identifies optimal flux patterns under given constraints and objectives, making it well-suited to reveal how anatomical organization and environmental conditions shape emergent metabolic states. Initial model analyses revealed that growth rates were identical across all simulations due to carbon limitation, while flux patterns varied depending on anatomical constraints.

### Increasing BS:M volume ratio and plasmodesmatal connectivity are sufficient to quantitatively predict photosynthetic diversity along the C3-C4 spectrum

First, we investigated how changes in the BS-M anatomy affect the choice of photosynthetic type under ambient conditions, i.e. v_c_:v_o_ = 3:1. For BS-M anatomies between 0.5 and 2.2, we simulated metabolic fluxes in the two-cell-type models and classified the resulting flux patterns according to their corresponding photosynthetic type (Materials and Methods). Fig. 2A shows representative flux maps obtained with BS-M anatomies of 0.5, 1.5, 1.8 and 2 which correspond to C3, C3-C4 type I, C3-C4 type II and C4 photosynthesis. The flux-based classifications closely match their corresponding anatomical categorisations, i.e., the model generally reproduces the photosynthetic types of the species with known volume ratio approximations. This supports the validity of our modelling approach.

**Figure 2:**
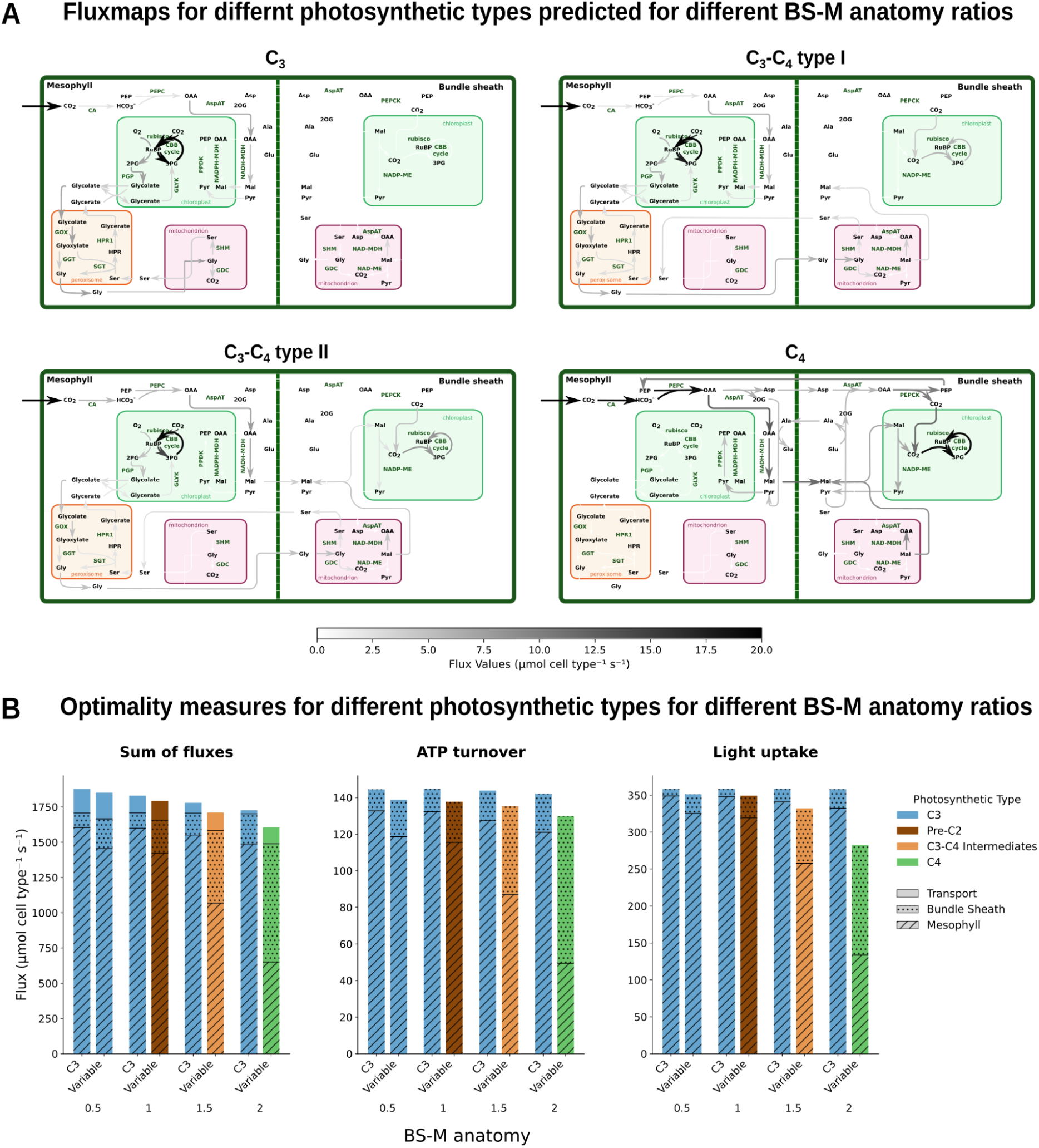
Predicted photosynthetic types and optimality measures for different BS-M anatomies at a rubisco carboxylation-to-oxygenation ratio of 3:1. A) Metabolic flux maps shown for four BS-M anatomies (0.5, 1.5, 1.8, and 2.0), representing C3, C3-C4 type I, C3-C4 type II, and C4 photosynthesis, respectively. Fluxes are reported per cell type. B) Sum of fluxes, ATP turnover, and light uptake (as a measure for quantum use efficiency) across different BS-M anatomies. For each BS-M anatomy value the left column shows values when C3 photosynthesis was enforced (by blocking bundle sheath rubisco) and the right column shows values for the predicted optimal photosynthetic type. The contributions of mesophyll cells, bundle sheath cells, and transport reactions to each optimality measure are shown as striped, dotted areas and plain, respectively. Fluxes per cell type are summed and reported per m^-2^ leaf. Abbreviations: CBB cycle - Calvin-Benson-Bassham Cycle, Reactions: AspAT - aspartate transaminase (EC 2.6.1.1), CA - carbonic anhydrase (EC 4.2.1.1), GDC - glycine decarboxylase (EC 1.4.1.27), GGT - Glutamate-glyoxylate aminotransferase (EC 2.6.1.4), GLYK - Glycerate kinase (2.7.1.31), GOX - glycolate oxidase (EC 1.1.3.15), HPR1 - hydroxypyruvate reductase (EC 1.1.1.29), NAD-MDH - NAD-malate dehydrogenase (EC 1.1.1.37), NAD-ME - NAD-malic enzyme (EC 1.1.1.38), NADP-MDH - NADP-malate dehydrogenase (EC 1.1.1.82), NADP-ME - NADP-malic enzyme (EC 1.1.1.40), PEPC - PEP carboxylase (EC 4.1.1.31), PEPCK - PEP carboxykinase (EC 4.1.1.49), PGP - 2-phosphoglycolate phosphatase (EC 3.1.3.18), PPDK - Pyruvate phosphate dikinase (2.7.9.1), SGT - L-serine:glyoxylate aminotransferase (EC 2.6.1.45), SHM - serine hydroxymethylase (EC 2.1.2.1). Metabolites: 2PG - 2-phosphoglycolate, 3PG - 3-phospho-D-glycerate, Asp - aspartate, Gly - glycine, HPR - 3-hydroxypyruvate, Mal - malate, OAA - oxaloacetate, PEP - phosphoenolpyruvate, Pyr - pyruvate, Ser - serine.

In the C3 scenario, CO_2_ is fixed in the mesophyll by rubisco and due to the imposed photorespiratory constraints we observe an active photorespiratory pathway in the mesophyll. In the C3-C4 type I scenario, rubisco operates in both mesophyll and bundle sheath cells. Photorespiratory glycine produced in the mesophyll is transported to bundle sheath mitochondria, where glycine decarboxylase (GDC) releases CO_2_ for refixation by bundle sheath rubisco. Serine moves back to the mesophyll to complete the photorespiratory cycle. In the C3-C4 type II scenario, bundle sheath rubisco receives CO_2_ from both the glycine shuttle (as in type I) and a nascent C4 cycle. Phosphoenolpyruvate carboxylase (PEPC) fixes bicarbonate to oxaloacetate in the mesophyll, which is then reduced to malate and transported to the bundle sheath. There, plastidial NADP-malic enzyme (NADP-ME) decarboxylates malate, releasing CO_2_ and further increasing bundle sheath rubisco activity beyond type I levels. Finally, in the C4 scenario, rubisco activity is confined to bundle sheath cells and all mesophyll CO_2_ enters the C4 cycle via PEPC. Decarboxylation of the C4 acids malate and aspartate in the bundle sheath occur through combined activity of plastidial NADP-ME and cytosolic PEP carboxykinase (PEPCK). Alanine and PEP are shuttled back to the mesophyll, where alanine is transaminated to pyruvate, which is then phosphorylated to PEP by pyruvate phosphate dikinase (PPDK) thus completing the cycle.

Our model predictions relied on the assumption that maximal growth was achieved with minimal enzyme investment as represented by the minimization of reaction fluxes. To test the generalizability of this assumption we examined whether flux minimization would also translate into lower ATP turnover and higher quantum yield efficiency (as determined by the light uptake). We compared ATP turnover and light uptake of flux solutions obtained for different BS-M anatomies to corresponding solutions in which C3 photosynthesis was enforced and found a consistent pattern of decreasing flux sums and ATP turnover and increasing quantum yield efficiency as we transition from C3 to C4-typical BS-M anatomies. At the same time, we also observe a shift in the metabolic activity in terms of reaction flux, ATP turnover, and light uptake from the mesophyll to the bundle sheath (Fig 2B). These observations underline the generalizability of our conclusions independent of the assumed optimality criterion and reflect the spatial restructuring of carbon fixation during the C3 to C4 transition.

Overall, our model *ab initio* recapitulates different photosynthetic types through combined variation of the V_BS/M_ ratio and plasmodesmatal connectivity. These predictions quantitatively match anatomical expectations. These observations highlight the crucial role of leaf anatomy in enabling and shaping photosynthetic metabolism under constant ambient conditions.

### Changes in photorespiration and BS-M anatomy drive photosynthetic diversity and lead to the emergence of potential pre-C2 metabolic states

Next, we tested how elevated temperatures, as manifested in increased levels of photorespiration, affect the photosynthetic choice and leaf metabolism for different leaf anatomies. We varied photorespiration by changing the v_c_:v_o_-ratios of the mesophyll and bundle sheath rubisco populations. The bundle sheath rubisco population without oxygenase activity was left unaltered. Flux solutions were obtained along a gradient of increasing levels of photorespiration and BS-M anatomies and classified by their photosynthetic type (Materials and Methods) (Fig 3). Both, photorespiratory levels and BS-M anatomies drive photosynthetic type selection. Under low photorespiration and small BS-M anatomy values, flux solutions exhibit predominantly mesophyll rubisco activity and are classified as C3 photosynthesis. Conversely, high photorespiration and large BS-M anatomy values yield predominantly bundle sheath rubisco activity and classify as C4 photosynthesis. C3-C4 intermediates emerge under ambient and near-ambient photorespiratory levels and intermediate BS-M anatomies. Thus, our model successfully recapitulates the transitory steps from C3 to C4 photosynthesis along a combined gradient of changes to leaf anatomy and environmental conditions.

**Figure 3:**
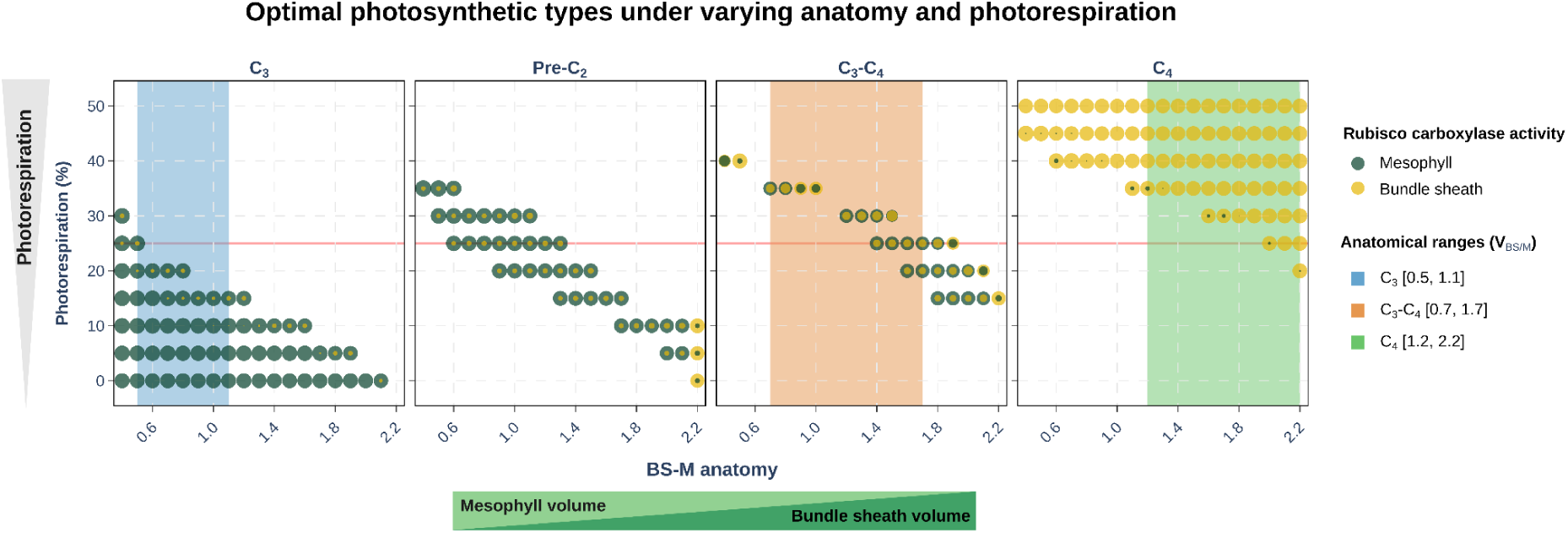
Predicted photosynthetic types for different BS-M anatomies and levels of photorespiration. Flux solutions were classified by photosynthetic type and displayed in separate panels. Each bubble represents an individual flux solution. Bubble colours indicate rubisco activity in the mesophyll or bundle sheath and bubble size encodes rubisco flux value. Overlapping bubbles indicate concurrent activity of both reactions. The red line marks the rubisco v_c_/v_O_ ratio of 3:1 assumed under ambient conditions. Coloured backgrounds indicate the anatomically observed V_BS/M_-ratio ranges for each photosynthetic type, derived from published data (see Supplementary Information).

To dissect the contribution of plasmodesmatal flux weighting to the flux predictions we varied V_BS/M_ only and did not assign any anatomy-dependent weights to the plasmodesmatal fluxes. In these simulations the model reproduced the C3-C4 transitions, however, it favoured C4 photosynthesis even under conditions where it would not be expected to occur, namely, low photorespiratory conditions and C3-typical values of V_BS/M_ (Supplementary Fig. S4). This underlines that the tradeoff in metabolic cost between operating photorespiration or a C4 cycle is not sufficient to explain the known ecology of C3 and C4 plants and that additional constraints which limit efficient metabolite transport operate *in vivo*. Together, this highlights that concerted changes to cell type metabolic capacity and connectivity efficiently drive the C3-C4 transition.

Notably, flux solutions in the transition zone between C3 and C3-C4 classifications included states that did not match any previously described photosynthetic type. We refer to them here as “pre-C2”. These flux solutions display bundle sheath rubisco activity using CO_2_ from decarboxylation reactions, however, via non-canonical enzymes (not GDC, PEPCK, NADP-ME, or NAD-ME). These enzymes vary across different flux solutions and result in a total of 16 decarboxylation reactions which group into amino acid biosynthesis, fatty acid biosynthesis, and central carbon metabolism pathways. Thermodynamic analysis confirmed the thermodynamic feasibility of all predicted pre-C2 decarboxylation reactions through negative ΔG′° values (Materials and Methods and Supplementary Table S7).

To further investigate these non-canonical CO_2_-concentrating pathways we inspected those reactions that appeared in at least 80% of all pre-C2 flux solutions and contributed more than 5% to the total flux. These reactions are plastidial acetolactate synthase (ASL, EC 2.2.1.6) and β-isopropylmalate dehydrogenase (IPMDH, EC 1.1.1.85) from branched-chain amino acid biosynthesis, mitochondrial pyruvate dehydrogenase (PDH, EC 1.2.1.104) and cytosolic isocitrate dehydrogenase (ICDH, EC 1.1.1.42) (Fig 4). We observe two main decarboxylation pathways which are fueled by some partial C4 cycle activity involving PEPC and malate and Asp synthesis. Asp and malate are shuttled to the bundle sheath cells. Asp is converted to malate and malate and pyruvate fuel the bundle sheath TCA cycle driving the decarboxylation reactions of mitochondrial PDH and the synthesis of citrate. Part of the generated citrate is transported to the cytosol and decarboxylated by ICDH. Another part of the bundle sheath pyruvate is used in the bundle sheath plastid for leucine and valine biosynthesis involving decarboxylation steps via ALS and IPMDH. Investigating cell-type specific nitrogen assimilation across C3, pre-C2, C3-C4, and C4 solutions reveals that nitrate and nitrite reduction is mesophyll-confined in strict C3 but occurs in both cell types across other solutions. Glutamine synthase activity shows a progressive mesophyll-to-bundle-sheath shift, with considerable flexibility within photosynthetic types (Supplementary Fig. S5).

**Figure 4:**
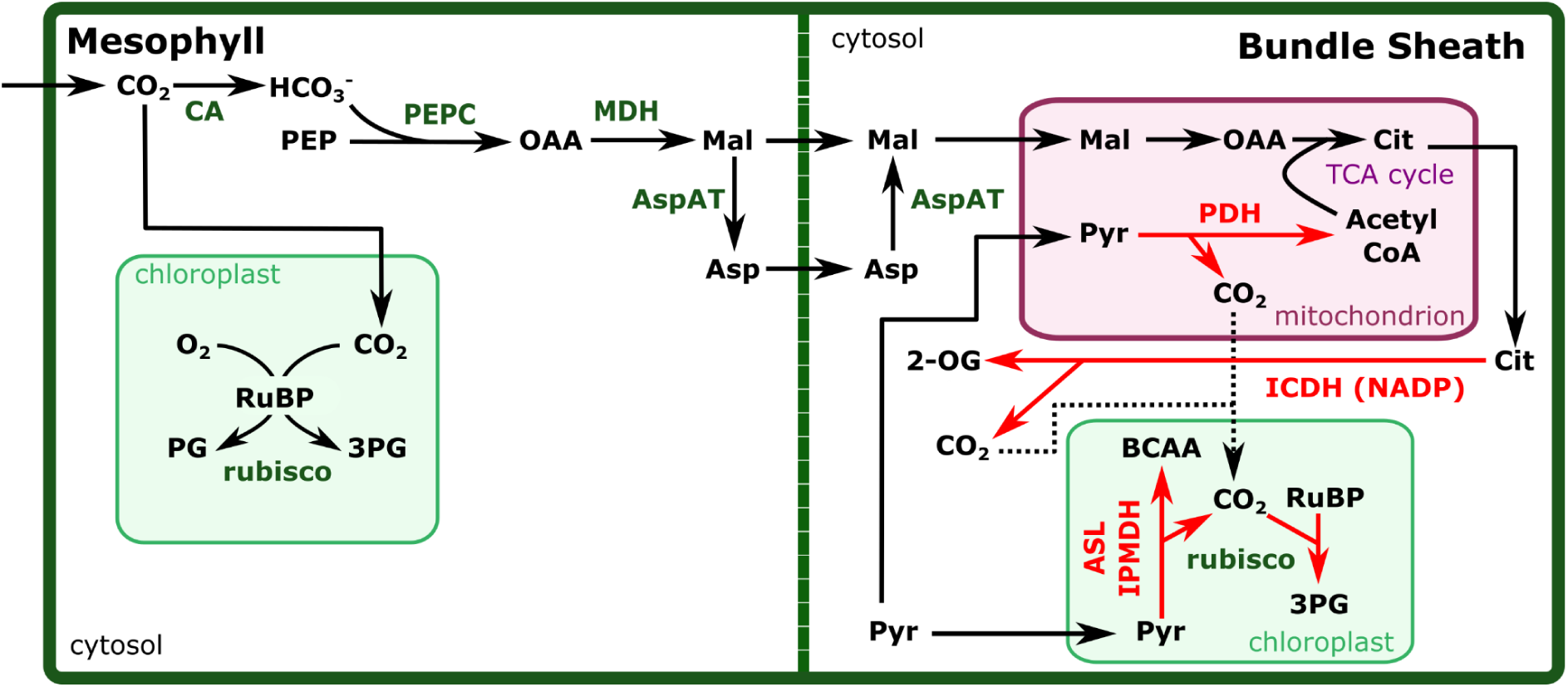
Fluxmap of pre-C2 flux modes. Flux modes supporting four non-canonical CO_2_-releasing reactions in the bundle sheath present in at least 80% of the pre-C2 flux solutions and contributing more than 5% to the total flux. Reaction names are present in green. Red arrows represent non-canonical CO_2_-releasing reactions. Abbreviations: Reactions: ASL - acetolactate synthase (EC 2.2.1.6), AspAT - aspartate transaminase (EC 2.6.1.1), CA - carbonic anhydrase (EC 4.2.1.1), ICDH - isocitrate dehydrogenase (EC 1.1.1.42), IPMDH - β-isopropylmalate dehydrogenase (EC 1.1.1.85), MDH - malate dehydrogenase (EC 1.1.1.37), PDH - pyruvate dehydrogenase (EC 1.2.1.104), PEPC - PEP carboxylase (EC 4.1.1.31); Metabolites: 2-OG - 2-oxoglutarate, 3PG - 3-phospho-D-glycerate, Asp - aspartate, BCAA - branched chain amino acids (leucine and valine), Cit - citrate, Mal - malate, OAA - oxaloacetate, PEP - phosphoenolpyruvate,Pyr - pyruvate.

Overall, this suggests that concurrent activity of rubisco and PEPC in the mesophyll together with a shift in branched-chain amino acid metabolism toward the bundle sheath cells and increased bundle sheath cell TCA cycle activity can provide energetic advantages under conditions of elevated photorespiration.

### Accounting for bundle sheath cell wall suberisation predicts C4 grasses photosynthetic metabolism

Many C4 grasses possess suberised bundle sheath cell walls, which limit CO_2_ leakage back into the mesophyll (Hatch 1987; Mertz and Brutnell 2014). This anatomical specialisation is typically accompanied by a reduction or loss of PSII activity and a dominant engagement of the NADP-ME subtype (Rao and Dixon 2016). The loss of PSII shifts the balance of electron transport away from linear electron transport (LET) towards cyclic electron transport (CET) (Oswald et al. 1990; Sheen et al. 1987). This shift has two complementary consequences. First, reduced PSII activity decreases O_2_ evolution from water-splitting at PSII in the bundle sheath chloroplast, thereby minimising local photorespiration and enhancing the efficiency of the carbon-concentrating mechanism (Maroco et al. 1998). Second, it has been shown that the shift towards CET helps balance the ATP/NADPH ratio to meet the higher ATP demands of C4 photosynthesis (Takabayashi et al. 2005). Since CET generates only ATP whereas LET produces both ATP and NADPH at a ratio of ∼1.3 (Foyer et al. 2012), decreasing LET raises ATP output without a corresponding increase in NADPH production. By engaging the NADPH-ME subtype, decarboxylation of malate in the bundle sheath chloroplast directly supplies NADPH from the mesophyll, thus compensating for the downregulation of PSII (Hatch 1987).

To assess the effect of suppressed bundle sheath leakage in our model, we simulated suberised bundle sheath cells by blocking CO_2_ and O_2_ exchange between the two cell types and compared this to the previous scenario with non-suberised bundle sheath cells, where gases were allowed to be freely exchanged.

We first compared the two C4 scenarios (BS-M anatomy = 2) and focused on the activity of PSI and PSII and the contributions of LET and CET to energy and reducing equivalent provision (represented in Fig 5A). We quantified LET as the flux through ferredoxin-NADP(+) oxidoreductase (FNR) (EC 1.18.1.2) and CET as the flux through the photosynthetic NAD(P)H dehydrogenase (NDH) complex (EC 7.1.1.-). In the non-suberised scenario, both photosystems were highly active in the bundle sheath at a PSI:PSII ratio of ∼ 4.5 which translates into high levels of LET and a smaller contribution of CET at a ratio of LET:CET of ∼ 6.2 (Fig 5B). In contrast to this, when the bundle sheath is suberised, both photosystems show markedly reduced activity, with PSI flux declining to ∼ 46 % and PSII flux dropping to just ∼ 20% of non-suberised levels. As a consequence bundle sheath electron transport is shifted towards higher CET and results in a markedly lower LET:CET ratio of ∼0.62 when compared to the non-suberised scenario. Concurrently, in the suberised scenario NADP-ME emerges as the primary decarboxylation enzyme with a minor contribution from PEPCK, in contrast to the non-suberised scenario where PEPCK predominates alongside NADP-ME (Fig 5C).

**Figure 5:**
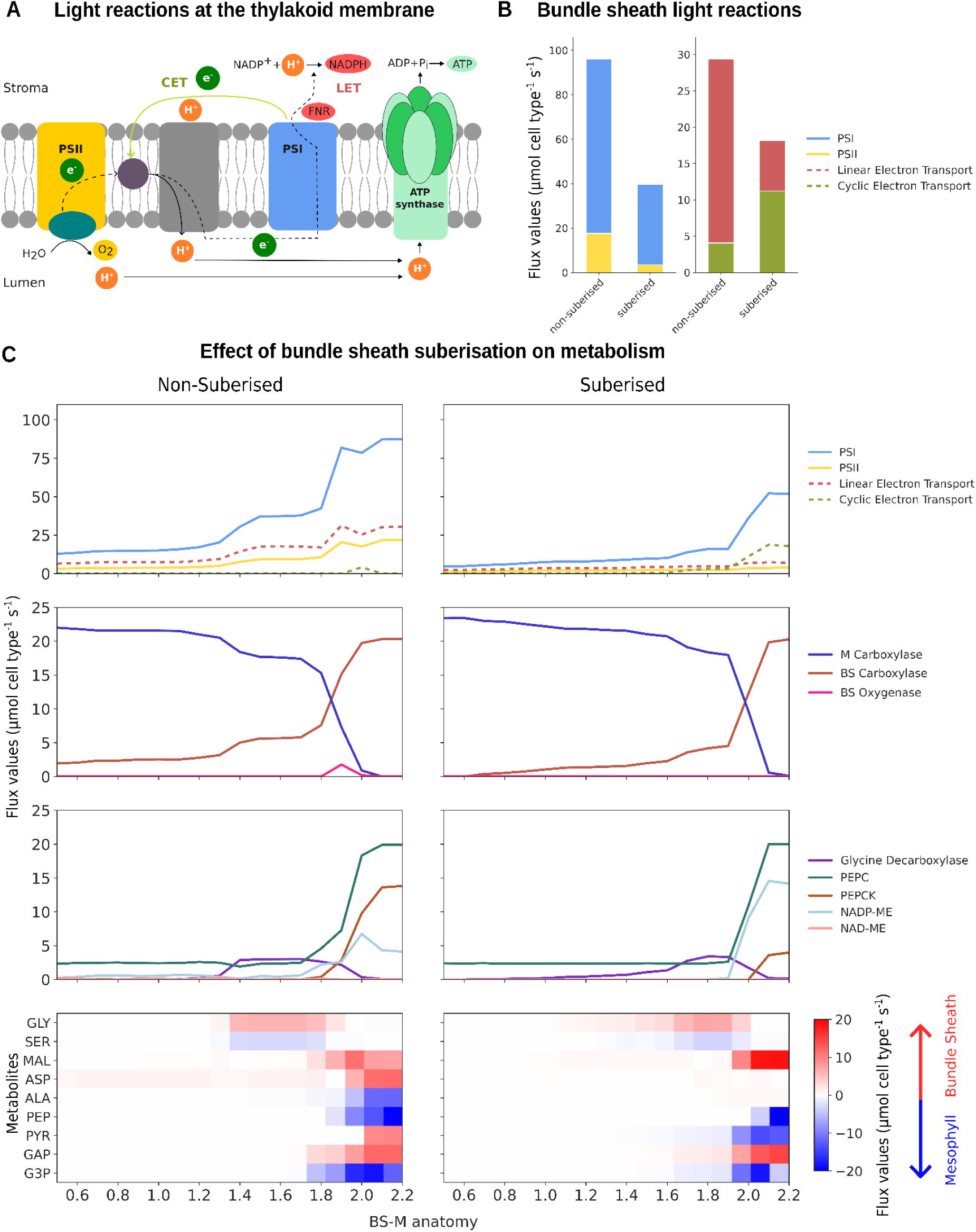
Effect of suberisation and bundle sheath to mesophyll volume ratio on photosynthesis and metabolism. A) Schematic representation of the thylakoid membrane electron transport chain. B) Light reactions in C4 photosynthesis (BS-M anatomy = 2) for non-suberised and suberised bundle sheath cell walls. C) Effects of cell wall suberisation on photosystem activities, linear and cyclic electron transport, photorespiration, decarboxylation enzyme selection, and metabolite exchange between cell types across the C3–C4 spectrum. In the heatmaps, mesophyll-to-bundle sheath transport fluxes are shown in red; bundle sheath-to-mesophyll fluxes are shown in blue. Flux values are reported per cell type. Abbreviations: ALA - alanine, ASP - aspartate, CET - cyclic electron transport, FNR - ferredoxin-NADP(+) oxidoreductase, G3P - 3-phosphoglycerate, GAP - 3-phosphoglyceraldehyde, GLY - glycine, LET - linear electron transport, MAL - malate, PEP - phosphoenol-pyruvate, PYR - pyruvate.

Thus, our model analyses suggest that when O_2_ is trapped inside the bundle sheath due to cell wall suberisation, a reduction in PSII is required to not compromise the established carbon concentration mechanism (CCM) by accumulating O_2_. The resulting shift from LET to CET and from NADPH and ATP production towards exclusive ATP production is compensated by the NADPH supply from the mesophyll via NADP-ME.

These observations prompted us to conduct a system-wide analysis of the effects of bundle sheath cell wall suberisation on photosystem activities and LET and CET, photorespiration, the choice of decarboxylation enzymes, and metabolite exchange between cell types across the C3-C4 spectrum (Fig 5C). When comparing non-suberised and suberised conditions, we observe a consistently higher engagement of bundle sheath PSI and PSII in the non-suberised scenario. In the suberised scenario, PSII is continuously suppressed across the C3-C4 spectrum, and from an intermediate BS-M anatomy of ∼1.6 onwards CET is dominant (Fig 5C top row). The low basal level of O_2_ evolving at PSII under suberised conditions is used as an electron acceptor for mitochondrial respiration, such that photorespiration is absent in the bundle sheath across all BS-M anatomies. In contrast to this, for the non-suberised scenario, we observe some photorespiration at intermediate levels when diffusion and decarboxylation events through GDC and the PEPC-driven C4 cycle provide CO_2_ (Fig. 5C second row). As in the C4 scenario, PEPCK emerges as the main decarboxylation enzyme, supported by substantial NADP-ME flux in the non-suberised scenario, whereas NADP-ME is the primary decarboxylation enzyme, with smaller PEPCK contributions in the suberised scenario. We see NAD-ME to only have very low contribution in the non-suberised scenario (Fig. 5 third row).

Finally, we observe differences in the metabolites shuttled between the mesophyll and the bundle sheath (Fig 5 bottom row). At intermediate BS-M anatomies glycine transport to the bundle sheath for the GDC is similar in both scenarios. However, at C4 BS-M anatomies, both ASP and MAL are shuttled into the bundle sheath in the non-suberised scenario, consistent with the co-dominance of PEPCK and NADP-ME, whereas MAL is the predominant C4 acid in the suberised scenario, reflecting the dominance of NADP-ME. Correspondingly, PEP and ALA are the main return metabolites to the mesophyll in the non-suberised scenario, while PYR and PEP fulfil this role in the suberised scenario. Both patterns are consistent with the known biochemistry of the respective C4 subtypes (Weber and von Caemmerer 2010; Rao and Dixon 2016; Arrivault et al. 2017). Lastly, the triose-phosphate shuttle transfers NADPH and ATP between the mesophyll and bundle sheath cells to balance the energy demands in both scenarios and particularly in the NADP-ME subtype where bundle sheath PSII activity is reduced (Arrivault et al. 2017). In accordance with this, our model predicts the emergence of this shuttle for C4 BS-M anatomies in both scenarios with a slightly higher flux in the suberised scenario.

In summary, these observations emphasize the structure-function relationship on the C3-C4 spectrum and highlight the role of bundle sheath suberisation in shaping photosynthetic metabolism.

### C3-C4 intermediates provide an energetic advantage under high bundle sheath leakage and photorespiration

In our analysis we assume that suberised bundle sheath cells completely lack gas exchange between the mesophyll and the bundle sheath. In reality, however, even a fully impermeable cell wall would permit some gas diffusion via plasmodesmata. We therefore investigated how varying degrees of CO_2_ leakage from the bundle sheath, in combination with different photorespiratory conditions, influence energetic optimality across the C3-C4 spectrum.

We implemented leakage by enforcing CO_2_ export from the bundle sheath as a percentage of all CO_2_ transported into or generated in the bundle sheath. Except for the fully suberised scenario as an extreme case, O_2_ diffusion to or from the bundle sheath was not constrained because we do not have knowledge about the actual concentration gradients. We predicted metabolic flux patterns for a BS-M anatomy of 1.5 (resembling C3-C4 anatomy) and classified them as C3, pre-C2, C3-C4 intermediates or C4 photosynthesis. If a flux pattern was classified as pre-C2 or C3-C4 intermediate, its energetic efficiency, measured by ATP turnover, was compared against two reference scenarios: a C3 scenario in which rubisco activity was confined to the mesophyll, and a C4 scenario in which rubisco activity was confined to the bundle sheath. The reduction in ATP turnover relative to the second most efficient solution was then plotted. For flux patterns classified as C3 or C4, the reduction relative to the other reference scenario was plotted (Fig. 6).

**Figure 6:**
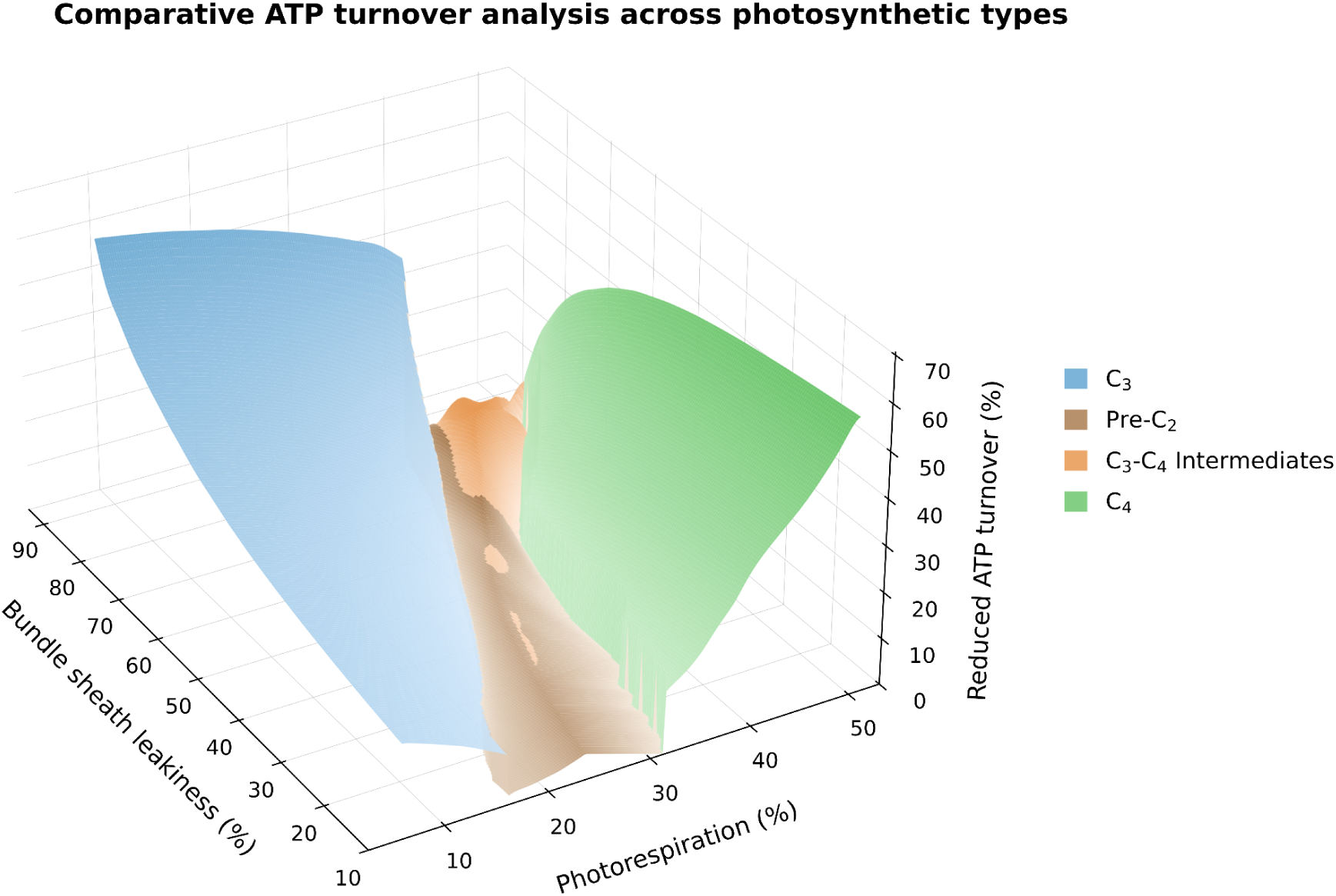
Photosynthetic optimality across bundle sheath leakage and photorespiration for C3-C4 anatomy (BS-M anatomy = 1.5). Metabolic flux patterns were classified as C3, pre-C2, C3-C4 intermediate, or C4 photosynthesis. For flux patterns classified as C3-C4 intermediates, energetic efficiency, as measured by ATP turnover, was benchmarked against two reference scenarios: a C3 scenario confining rubisco activity to the mesophyll, and a C4 scenario confining rubisco activity to the bundle sheath. The reduction in ATP turnover relative to the second most efficient reference scenario was then calculated and plotted. For flux patterns classified as C3 or C4, the reduction relative to the other reference scenario was plotted.

When bundle sheath cell leakage is high and photorespiration is low C3 photosynthesis is most energy efficient. Conversely, when bundle sheath cell leakage is low and photorespiration is high C4 photosynthesis is most energy efficient. Interestingly, between these two opposing optima, a corridor emerges along the trajectory of increasing bundle sheath cell leakage and photorespiration, in which intermediate photosynthetic types (pre-C2 or C3-C4 intermediates) represent a local energetic optimum. It shall be noted that the energetic advantage of intermediate photosynthesis reached a maximum of ∼15% while the energetic advantages of C3 and C4 over the second best reference scenario can reach up to ∼ 74 and ∼59%, respectively, for the simulated set of parameters.

We repeated the analysis for BS-M anatomies of 0.5., 1, and 2 and find this local optimum to be broader and higher as BS-M anatomy values increase, suggesting that C3-C4 intermediacy can represent a stable evolutionary state whose energetic advantage is shaped by the interplay of bundle sheath leakage, photorespiratory conditions, and BS-M anatomies (Supplementary Fig. S6). Conversely, this also implies that the evolutionary transition to C4 photosynthesis requires not only increased plasmodesmatal connectivity and bundle sheath cell volume, but also sufficiently low bundle sheath cell leakage.

## Discussion

Using an anatomy-aware flux-balance model of mesophyll and bundle sheath metabolism, we explored structure–function relationships across the C3-C4 spectrum. By varying anatomical parameters — including the V_BS/M_-ratio and plasmodesmatal connectivity — the model successfully recapitulates key photosynthetic types along this spectrum. Our analysis leads to four central findings:

### Metabolic cost alone does not explain photosynthetic diversity

When plasmodesmatal fluxes were not weighted by anatomy, the C3-to-C4 transition occurred with increasing V_BS/M_-ratios. However, C4 photosynthesis was favoured even under low photorespiratory conditions and C3-typical V_BS/M_, indicating that differences in metabolic cost alone do not explain the known distribution of C3 and C4 plants and that other limiting factors are involved. Plasmodesmatal connectivity emerged as a main enabler of the transition, determined by tight mesophyll-to-bundle-sheath contact and high plasmodesmatal density. This is consistent with empirical data, as C4 leaves contain up to ten times more plasmodesmata at this interface than C3 leaves (Danila et al. 2016, Schreier et al. 2022, Aleksejeva and Schreier 2025). It also agrees with the anatomical enabler and preconditioning hypotheses, according to which C4 origins are concentrated in lineages whose C3 members already possess enlarged bundle sheaths which enable tight mesophyll-to-bundle-sheath contact (Sage 2004, Christin et al. 2013, Griffiths et al. 2013). Increased bundle-sheath size and vein density (the latter cannot readily be represented in a constraint-based framework) are the principal quantitative anatomical changes (McKown and Dengler 2007, Lundgren et al. 2014) and proto-Kranz and C2 anatomy link these shifts to a photorespiratory CO_2_ pump (Sage et al. 2014).

Notably, increasing plasmodesmatal density for efficient metabolite exchange and decreasing bundle sheath cell leakage are two competing objectives that shape a conditions-dependent Pareto frontier (Kromdijk et al. 2014). The quantitative significance of this trade-off is underscored by estimates that CO_2_ diffusion via plasmodesmata accounts for up to 60% of total bundle sheath leakage in C4 species (Jenkins et al. 1989). CO_2_-diffusion via plasmodesmata represents the dominant route of bundle sheath leakage in suberised C4 species (Weiner et al. 1988; Danila et al. 2016; Alonso-Cantabrana et al. 2018). Yet despite its importance, the developmental mechanisms governing plasmodesmatal density, including the genetic regulators of plasmodesmata biogenesis and the direct impact of plasmodesmatal morphology on photosynthetic performance, are still poorly understood (Schreier et al. 2026). Quantitative studies of this trade-off using more anatomy-explicit models shall guide engineering strategies for optimized cell-to-cell connectivity in crops as more of the underlying regulatory networks are being characterised.

Moreover, experimental and computational studies have suggested that C4 photosynthesis is less phenotypically plastic than C3 photosynthesis, which may contribute to the more ecologically restricted occurrence of C4 (Sage and McKown 2006; Sundermann et al. 2021). In contrast, many C3-C4 intermediates engage in facultative use of the photorespiratory CO_2_ pump which holds the potential for greater phenotypic plasticity (Schlüter et al. 2023, Leung et al. 2026). Our analysis supports this view: intermediates span the largest physiologically relevant region of the investigated parameter space across BS-M anatomies and photorespiratory levels, covering a continuum from very weak to strong C4-like character. This has implications for crop engineering. Intermediates occupy a region of parameter space that is potentially accessible through reversible metabolic adjustments. For crops targeted at warmer, more variable climates, a quantitative analysis of growth benefits across anticipated environmental scenarios could therefore inform whether engineering toward an intermediate, rather than a full C4 state, represents the more robust and achievable strategy.

### Shifting branched-chain amino acid metabolism and TCA cycle activity towards the bundle sheath acts as a non-canonical carbon-concentrating mechanism

Our analysis identifies potential pre-C2 metabolic states in which concurrent mesophyll rubisco and PEPC activity, together with bundle sheath-directed branched-chain amino acid metabolism and TCA cycle activity, confer energetic advantages under elevated photorespiration. Notably, predicted PEPC activity followed a non-monotonic pattern with increasing values of BS-M anatomies. Activity was initially elevated at low BS-M anatomy values, transiently reduced upon the onset of GDC activity, and subsequently rose sharply with the establishment of the full C4 cycle (Fig 5C). The joint activity of rubisco and PEPC in the mesophyll is a defining feature of early C3-C4 evolution (Gowik and Westhoff 2011).

Beyond carbon metabolism, our model predicts both mesophyll- and bundle sheath-localised nitrogen assimilation as optimal states across pre-C2 solutions (Supplementary Figure S4). This dual localisation reflects a broader body of experimental evidence suggesting that bundle sheath cells in C3 leaves can harbour a specialised metabolism, which may underpin the pre-C2 states reported here. In Arabidopsis, for instance, glutamate synthases and glutamate dehydrogenase mRNAs were shown to be abundant on bundle sheath ribosomes (Aubry et al. 2014) and reticulate mutants revealed that a number of enzymes of amino acid metabolism are enriched in the vasculature and bundle sheath (Rosar et al. 2012; Lundquist et al. 2014). A study in rice demonstrated that transcripts for nitrate and nitrite reduction were more highly expressed in bundle sheath and veinal cells than in mesophyll cells, while glutamine and glutamate synthase were expressed in both cell types - indicating that the bundle sheath is specifically tuned for nitrate assimilation and amino acid biosynthesis (Hua et al. 2021). The degree of this compartmentation is not fixed. Work in rice and barley has shown that the cellular partitioning of ammonium assimilation is strongly influenced by nitrogen nutritional status (Tobin and Yamaya 2001).

Altogether, pre-C2 flux modes identified by our model, represent a state with partial CO_2_ concentration without committed anatomical reorganisation which opens potential avenues for metabolic engineering. At the same time, our findings underscore the need for deeper, context-specific knowledge of bundle sheath cell metabolism in C3 plants - knowledge that is likely to differ not only among species but also across conditions (Leegood 2008).

### Minimization of bundle sheath O_2_ evolution is a key driver of PSII downregulation and NADP-ME dominance

Comparing non-suberised and suberised C4 bundle sheath cell metabolism, suggest that when O_2_ is trapped inside the bundle sheath cells, a reduction in PSII is required to not compromise the established CCM by accumulating O_2_. This drives a shift from LET to CET and from NADPH and ATP production towards exclusive ATP production which is compensated by NADPH supply from the mesophyll via NADP-ME. Complementary to this, previous kinetic modeling has highlighted the need to keep the NADPH-to-NADP ratio low so that the driving force for NADP-ME decarboxylation is strong (Bräutigam et al. 2018). Although arguing from different starting points, both findings provide an explanation for the observed concurrent reduction in PSII and dominance of the NADP-ME subtype. Beyond suberisation, additional metabolic constraints not considered here likely contribute, including those related to C4 acid import into bundle sheath chloroplasts (Arrivault et al. 2025). A rich body of literature has discussed the potential environments and physico-chemical constraints that led to the evolution of 3 different enzymes as main decarboxylation enzymes (Blätke and Bräutigam 2019; Hatch 1987; Sage 2004; Furbank 2011) and our model provides some mechanistic underpinning for the observed co-occurrence of bundle sheath suberisation, loss of PSII and NADP-ME engagement in some C4 grasses.

### Energetic advantage of C3-C4 intermediates supports the notion of intermediate photosynthesis as a stable evolutionary state

Our results suggest that intermediate photosynthesis represents a local optimum along the anatomical gradient toward C4, conferring improved photosynthetic performance under specific environmental conditions without requiring the full biochemical and anatomical remodelling of C4 plants. While photorespiration is a well-studied driver of C3-C4 and C4 evolution, the role of bundle sheath leakage has received comparatively little attention. Our analysis highlights leakage not only as a determinant of C4 efficiency but as a key factor governing the productivity gains of intermediate photosynthesis over both C3 and C4 biochemistries.

Reduced leakage can be achieved through several mechanisms operating at different spatial scales. At the cell wall level, suberisation provides a fixed physical barrier to CO_2_ diffusion, tightly coupled to PSII reduction in full C4 monocots, though its extent in C3-C4 intermediates remains largely unknown. At the subcellular level, organelle repositioning offers an additional means of modulating effective leakage, and notably occurs prior to the establishment of the glycine shuttle during proto-Kranz and pre-C2 anatomy (Lundgren 2020; Sage et al. 2012). Since our model operates at the cellular level, these determinants - including differential chloroplast and mitochondria numbers and positioning in mesophyll and bundle sheath cells - are subsumed into a single leakage term. Nevertheless, even this aggregate treatment highlights leakage as a quantitatively important parameter, underscoring the need for more quantitative experimental and modelling work to dissect its individual cellular and subcellular contributors.

### Limitations of the model and future perspectives

Although our model is quantitative, predictions need to be treated in a semi-quantitative manner as our approach to divide the leaf into mesophyll and bundle sheath cell-type enriched volumes can only serve as a first proxy to capturing much more complex anatomical features. Besides subcellular organization which should be accounted for in future analyses, other, more specialised anatomical features should receive more attention. For instance, some C4 grasses have evolved specialized cells, the so-called bundle sheath extensions, which act as channels that optimize light availability to the bundle sheath and mesophyll (Bellasio and Lundgren, 2016 and Karabourniotis et al., 2000). Also the role of vasculature and reduced interveinal distances in the C3 to C4 transition is not accounted for (Schlüter and Weber 2020). Novel 3D leaf imaging techniques can provide the data to parameterize more fine-grained predictive models and will allow us to study the inner reality of leaves with high resolution and accuracy (He et al. 2025; Xiao et al. 2023; Retta et al. 2023). This will also enable us to understand species and condition-specific leaf trait characteristics and how this interplays with photosynthetic performance.

Our findings highlighted the role of plasmodesmatal connectivity and bundle sheath cell leakage as important factors in shaping photosynthetic metabolism. Together with intercellular gas and metabolite concentration gradients, these anatomical traits determine diffusional exchange fluxes and the efficiency of a bundle sheath cell CCM. As in any stoichiometry-based flux modelling approach our model does not explicitly account for metabolite concentrations and diffusional processes. Instead, we relied on an implicit representation by adding a decreasing penalty to exchange fluxes which mimic the increasing symplastic conductance along the C3-C4 spectrum and by enforcing a fixed percentage of CO_2_ that was transported to the bundle sheath cells to be exported. Further, as the volume of mesophyll intercellular air spaces is reduced in C4 compared to C3, mesophyll conductance decreases and potentially limits rubisco-based primary C-fixation, which generally has lower catalytic efficiency compared to PEPC and its substrate bicarbonate. Future studies shall explicitly account for diffusion processes to resolve those underlying constraints.

Taken together, our study reveals the intricate interplay between leaf anatomy, metabolism, and environment across the C3-C4 spectrum, and highlights the unresolved question of whether intermediate C3-C4 metabolism is shaped by, or itself shapes, the anatomical transitions that accompany C4 evolution.

## Material and Methods

### Derivation of BS: M volume ratios

We approximate the BS and M cell-enriched volumes as cuboids, calculated as the product of each cell-type enriched volume’s width (*W_BS_* and *W_M_*), length (*L_BS_*and *L_M_*), and thickness (*D_BS_* and D*_M_*) of the leaf. We assumed that the parameters length and thickness of the cuboids to be equal for the M and BS cell type-enriched volumes, which simplifies the equation for the BS:M volume ratios (V_BS/M_) to the area ratio (A_BS/M_), and ultimately to the width ratio of *W_BS_* and *W_M_*.

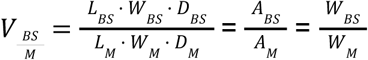

Note, that the assumption that the volume ratio of BS:M is equal to the area ratio of BS:M is important as fluxes are given per m^2^.

### V_BS/M_-dependent weighting of plasmodesmatal fluxes (α)

Metabolite exchange between M and BS is constrained by two key anatomical features: cell-cell contact surface area and plasmodesma density (Danila et al. 2016), together termed here plasmodesmatal connectivity. To incorporate these physical constraints with our model, we introduced a mechanistic penalty factor α(*V_BS/M_*) that scales M-BS transport resistance as a continuous function of the bundle sheath volume fraction **φ**(*V_BS/M_*):

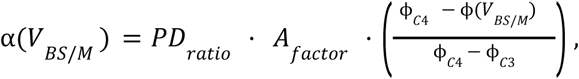

where PD_ratio_= 9.3 captures the fold-difference in plasmodesmatal frequency between fully developed C4 and baseline C3 species based on anatomical measurements from Danila et al. (2016). The contact surface area factor *Af_actor_* represents the proportional increase in BS-M interface:

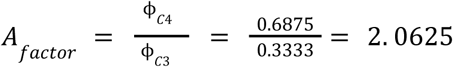

The term **φ**(*V_BS/M_*)represents the volume fraction as a function of the volume ratio *V_BS/M_*:

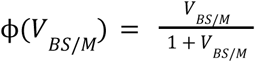

with **φ**_c3_ and **φ**_c4_ corresponding to the BS volume fractions associated with V_BS/M_ = 0.5 and V_BS/M_= 2.2, respectively.

### Weighted parsimonious flux balance analysis

We implemented weighted parsimonious flux balance analysis (wpFBA) by first solving an FBA problem in which growth, as represented by the flux through the biomass reaction, was maximized. Using this maximal biomass flux 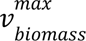 as a constraint, a second optimization then minimized the scaled and weighted sum of fluxes:

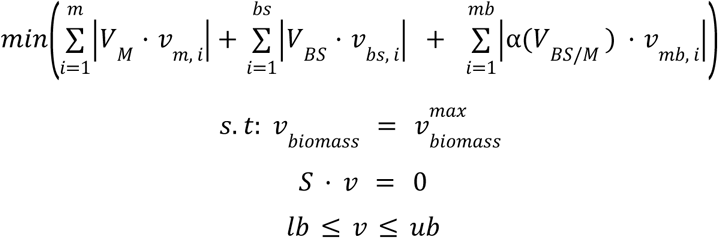

Fluxes in M and BS (as given in µmol m^-2^ s^-^) were scaled according to their cell type’s-enriched volume and exchange fluxes between M and BS were scaled according to the V_BS/M_-ratio (as detailed in SI, Anatomical constraints). Additionally, to model the increasing plasmodesmatal connectivity along the C3-C4 spectrum, plasmodesmatal transport fluxes between M and BS were weighted with a V_BS/M_-dependent penalty α.

The terms *m* and *bs* denote the total number of metabolic reactions in M and BS, respectively, and *mb* is the number of exchange reactions between the two cell types; 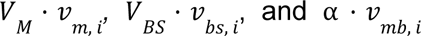 are the scaled fluxes in M, BS, and weighted, scaled exchange reactions between M and BS, respectively. *V_M_* and *V_BS_* are the M- and BS-enriched volumes, respectively. The vector *v* contains all reaction fluxes in the model, bounded by lower and upper flux boundaries *(lb)* and *(ub),* respectively.

### Implementation

Models were built and analysed using COBRApy (Ebrahim et al. 2013) in Python 3.7.10 using the solver Gurobi.

### Classifying flux solutions

Flux solutions were classified by photosynthetic type based on the presence or absence of flux through the reactions defined in Table 1. Reactions with scaled contributions of <10% of the total CO_2_ assimilation rate (< 2 µmol CO_2_ s^-1^) were excluded from the classification. For glycine decarboxylase, we used a threshold of <5% because it fixes two CO_2_ molecules per reaction event. Flux patterns emerging at BS-M anatomy values between C3 and C3–C4 type I were termed ‘pre-C2’. For C3–C4 type II and C4 classification, activity of at least one decarboxylation enzyme (PEPCK, NADP-ME, or NAD-ME) was required.

**Table 1:**
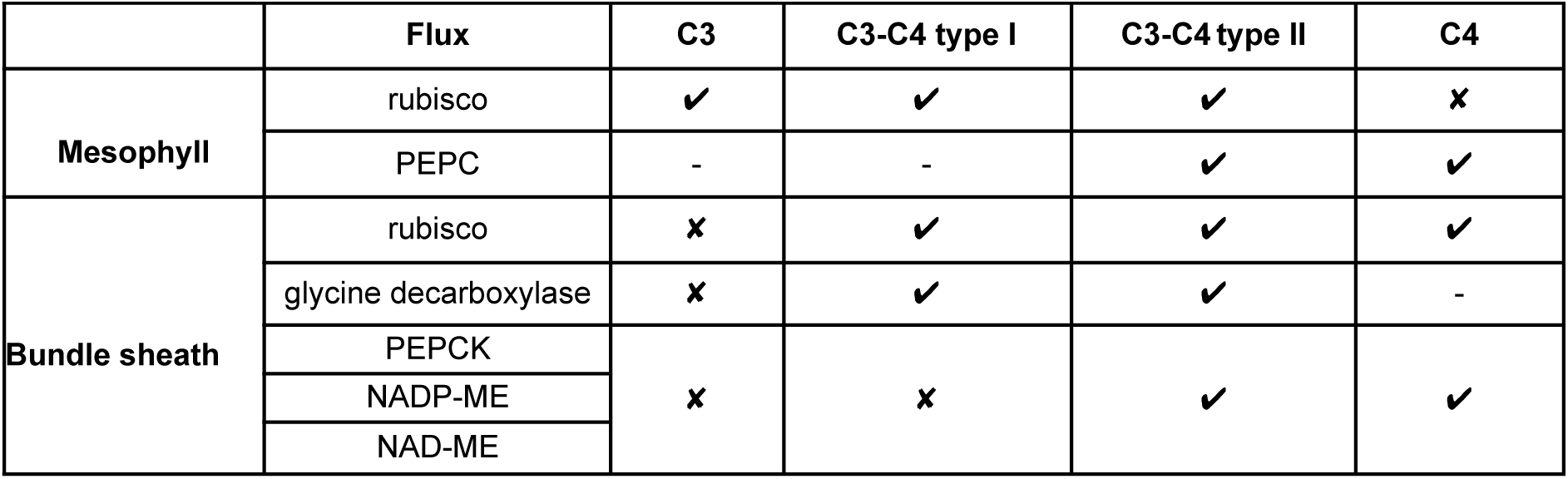
Classification of flux solutions by photosynthetic type. A checkmark indicates the presence of a defining flux, a cross indicates its absence, and a dash indicates that the reaction was not considered as a criterion for classification into that photosynthetic type.

### Thermodynamic analysis of non-canonical decarboxylation reactions

To evaluate the thermodynamic feasibility of the identified non-canonical pre-C2 decarboxylation reactions, we calculated standard transformed Gibbs free energy changes (ΔG′°) using values from AraCyc (Mueller et al. 2003) and independently verified these using eQuilibrator (Beber et al. 2022) (Supplementary Table S7). The use of standard conditions was necessitated by the absence of reliable intracellular metabolite concentration data. All reactions returned negative ΔG′° values, indicating thermodynamic feasibility, with the exception of isocitrate dehydrogenase. For this reaction, ΔG′° values of 1.57 kJ/mol (AraCyc) and 4.64 kJ/mol (eQuilibrator) are small and positive, suggesting that the reaction may be feasible under physiological conditions.

## Code availability

The code needed to reproduce the results of this paper is available at https://github.com/Toepfer-Lab/C3-C4_spectrum.

## Author Contribution

N.T. designed the research; A.L.-R. and T.M.M performed the research; S.D. and T.M.M collected and analyzed leaf anatomical data; T.M.M., A.L.-R., U.S., A.P.M.W. and N.T. analyzed data; T.M.M. drafted the article, N.T. wrote the article; T.M.M., A.L.-R. and N.T. created the figures.

## Acknowledgements

Figure 1 was created using BioRender under a CC-BY License.

## Bibliography

Aleksejeva, Kornelija, and Tina B. Schreier. 2025. “Stepwise Increase in Plasmodesmata During C4 Evolution in Flaveria.” bioRxiv. 10.1101/2025.04.18.649591.

Amthor, Jeffrey S., Arren Bar-Even, Andrew D. Hanson, et al. 2019. “Engineering Strategies to Boost Crop Productivity by Cutting Respiratory Carbon Loss.” The Plant Cell 31 (2): 297–314.

Arrivault, Stéphanie, David Barbosa Medeiros, Cristina Rodrigues Gabriel Sales, et al. 2025. “Metabolite Profiling Reveals Slow and Uncoordinated Adjustment of C4 Photosynthesis to Sudden Changes in Irradiance.” Plant Physiology 199 (3). 10.1093/plphys/kiaf508.

Arrivault, Stéphanie, Toshihiro Obata, Marek Szecówka, et al. 2017. “Metabolite Pools and Carbon Flow during C4 Photosynthesis in Maize: 13CO2 Labeling Kinetics and Cell Type Fractionation.” Journal of Experimental Botany 68 (2): 283–298.

Aubry, Sylvain, Richard D. Smith-Unna, Chris M. Boursnell, Stanislav Kopriva, and Julian M. Hibberd. 2014. “Transcript Residency on Ribosomes Reveals a Key Role for the Arabidopsis Thaliana Bundle Sheath in Sulfur and Glucosinolate Metabolism.” Plant J. 78 (4): 659–673.

Beber, Moritz E., Mattia G. Gollub, Dana Mozaffari, et al. 2022. “eQuilibrator 3.0: A Database Solution for Thermodynamic Constant Estimation.” Nucleic Acids Research 50 (D1): D603–D609.

Blätke, Mary-Ann, and Andrea Bräutigam. 2019. “Evolution of C4 Photosynthesis Predicted by Constraint-Based Modelling.” In bioRxiv. BioRxiv, June 13. 10.1101/670547.

Bräutigam, A., U. Schlüter, M. R. Lundgren, et al. 2018. “Biochemical Mechanisms Driving Rapid Fluxes in C 4 Photosynthesis.” In Plant Biology, No. Biorxiv;387431v1. BioRxiv, August 9. https://www.biorxiv.org/content/10.1101/387431v1.full.

Caemmerer, S. von. 2000. Biochemical Models of Leaf Photosynthesis. CSIRO Publishing.

Cheung, C. Y. Maurice, Mark G. Poolman, David A. Fell, R. George Ratcliffe, and Lee J. Sweetlove. 2014. “A Diel Flux Balance Model Captures Interactions between Light and Dark Metabolism during Day-Night Cycles in C3 and Crassulacean Acid Metabolism Leaves.” Plant Physiology 165 (2): 917–929.

Christin, P., C. P. Osborne, D. S. Chatelet, J. T. Columbus, G. Besnard, T. R. Hodkinson, L. M. Garrison, M. S. Vorontsova, and E. J. Edwards. 2013. “Anatomical enablers and the evolution of C4 photosynthesis in grasses.” Proc. Natl. Acad. Sci. U.S.A. 110 (4): 1381–1386.

Dal’Molin, Cristiana Gomes de Oliveira, Lake-Ee Quek, Robin William Palfreyman, Stevens Michael Brumbley, and Lars Keld Nielsen. 2010. “C4GEM, a Genome-Scale Metabolic Model to Study C4 Plant Metabolism.” Plant Physiology 154 (4): 1871–1885.

Danila, Florence R., William Paul Quick, Rosemary G. White, Robert T. Furbank, and Susanne von Caemmerer. 2016. “The Metabolite Pathway between Bundle Sheath and Mesophyll: Quantification of Plasmodesmata in Leaves of C3 and C4 Monocots.” The Plant Cell 28 (6): 1461–1471.

Danila, Florence R., William Paul Quick, Rosemary G. White, Steven Kelly, Susanne von Caemmerer, and Robert T. Furbank. 2018. “Multiple Mechanisms for Enhanced Plasmodesmata Density in Disparate Subtypes of C4 Grasses.” Journal of Experimental Botany 69 (5): 1135–1145.

Dusenge, Mirindi Eric, André Galvao Duarte, and Danielle A. Way. 2019. “Plant Carbon Metabolism and Climate Change: Elevated CO2 and Temperature Impacts on Photosynthesis, Photorespiration and Respiration.” New Phytologist 221 (1): 32–49.

Ebrahim, Ali, Joshua A. Lerman, Bernhard O. Palsson, and Daniel R. Hyduke. 2013. “COBRApy: COnstraints-Based Reconstruction and Analysis for Python.” BMC Systems Biology 7 (August): 74.

Foyer, Christine H., Jenny Neukermans, Guillaume Queval, Graham Noctor, and Jeremy Harbinson. 2012. “Photosynthetic Control of Electron Transport and the Regulation of Gene Expression.” Journal of Experimental Botany 63 (4): 1637–1661.

Furbank, Robert T. 2011. “Evolution of the C(4) Photosynthetic Mechanism: Are There Really Three C(4) Acid Decarboxylation Types?” Journal of Experimental Botany 62 (9): 3103–3108.

Gowik, Udo, and Peter Westhoff. 2011. “The Path from C3 to C4 Photosynthesis.” Plant Physiology 155 (1): 56–63.

Griffiths, H., G. Weller, L. F. M. Toy, and R. J. Dennis. 2013. “You’re so vein: bundle sheath physiology, phylogeny and evolution in C3 and C4 plants.” Plant, Cell & Environment 36: 249–261.

Gutteridge, Steven, and John Pierce. 2006. “A Unified Theory for the Basis of the Limitations of the Primary Reaction of Photosynthetic CO(2) Fixation: Was Dr. Pangloss Right?” Proceedings of the National Academy of Sciences of the United States of America 103 (19): 7203–7204.

Hatch, Marshall D. 1987. “C4 Photosynthesis: A Unique Elend of Modified Biochemistry, Anatomy and Ultrastructure.” Biochimica et Biophysica Acta 895 (2): 81–106.

Heckmann, David, Stefanie Schulze, Alisandra Denton, et al. 2013. “Predicting C4 Photosynthesis Evolution: Modular, Individually Adaptive Steps on a Mount Fuji Fitness Landscape.” Cell 153 (7): 1579–1588.

He, Jing, Kun Ning, Afroz Naznin, et al. 2025. “Technological Advances on Imaging and Modelling of Leaf Structural Traits: A Review on Heat Stress in Wheat.” Journal of Experimental Botany, ahead of print, March 4. 10.1093/jxb/eraf070.

Hua, Lei, Sean R. Stevenson, Ivan Reyna-Llorens, Haiyan Xiong, Stanislav Kopriva, and Julian M. Hibberd. 2021. “The Bundle Sheath of Rice Is Conditioned to Play an Active Role in Water Transport as Well as Sulfur Assimilation and Jasmonic Acid Synthesis.” The Plant Journal 107 (1): 268–286.

Jenkins, C. L. D., R. T. Furbank, and M. D. Hatch. 1989. “Mechanism of C4 Photosynthesis A Model Describing the Inorganic Carbon Pool in Bundle Sheath Cells.” Plant Phys 91: 1372–1381.

Kromdijk, Johannes, Nerea Ubierna, Asaph B. Cousins, and Howard Griffiths. 2014. “Bundle-Sheath Leakiness in C4 Photosynthesis: A Careful Balancing Act between CO2 Concentration and Assimilation.” Journal of Experimental Botany 65 (13): 3443–3457.

Langdale, Jane A. 2011. “C4 Cycles: Past, Present, and Future Research on C4 Photosynthesis.” The Plant Cell 23 (11): 3879–3892.

Leegood, Richard C. 2002. “C4 Photosynthesis: Principles of CO2 Concentration and Prospects for Its Introduction into C3 Plants.” Journal of Experimental Botany 53 (369): 581–590.

Leegood, Richard C. 2008. “Roles of the Bundle Sheath Cells in Leaves of C3 Plants.” Journal of Experimental Botany 59 (7): 1663–1673.

Leung, Arthur, Martha Ludwig, Tammy L. Sage, and Rowan F. Sage. 2026. “Acclimation to Enhanced Photorespiratory Conditions Increases the Effectiveness of C2 Photosynthesis in C3-C4 Intermediates.” bioRxiv. 10.64898/2026.09.22.753555.

Lundgren, Marjorie R., Colin P. Osborne, and Pascal-Antoine Christin. 2014. “Deconstructing Kranz Anatomy to Understand C4 Evolution.” Journal of Experimental Botany 65 (13): 3357–3369. 10.1093/jxb/eru186

Lundgren, Marjorie R. 2020. “C2 Photosynthesis: A Promising Route towards Crop Improvement?” New Phytologist 228 (6): 1734–1740.

Lundquist, Peter K., Christian Rosar, Andrea Bräutigam, and Andreas P. M. Weber. 2014. “Plastid Signals and the Bundle Sheath: Mesophyll Development in Reticulate Mutants.” Molecular Plant 7 (1): 14–29.

Machado, Tiago M., Nadine Töpfer, and Fatemeh Soltani. 2025. “Metabolic Modelling: Insights into the Machine Room of Plant Metabolism.” Journal of Plant Physiology 314 (154584): 154584.

Mallmann, Julia, David Heckmann, Andrea Bräutigam, et al. 2014. “The Role of Photorespiration during the Evolution of C4 Photosynthesis in the Genus Flaveria.” eLife 3 (June): e02478.

Maroco, J. P., Msb Ku, P. J. Lea, et al. 1998. “Oxygen Requirement and Inhibition of C4 Photosynthesis. An Analysis of c4 Plants Deficient in the c3 and c4 Cycles An Analysis of C4 Plants Deficient in the C3 and C4 Cycles.” Plant Physiology 116 (2): 823–832.

McKown, Athena D., and Nancy G. Dengler. 2007. “Key Innovations in the Evolution of Kranz Anatomy and C4 Vein Pattern in Flaveria (Asteraceae).” American Journal of Botany 94 (3): 382–399. 10.3732/ajb.94.3.382

Mertz, Rachel A., and Thomas P. Brutnell. 2014. “Bundle Sheath Suberization in Grass Leaves: Multiple Barriers to Characterization.” Journal of Experimental Botany 65 (13): 3371–3380.

Mueller, Lukas A., Peifen Zhang, and Seung Y. Rhee. 2003. “AraCyc: A Biochemical Pathway Database for Arabidopsis.” Plant Physiology 132 (2): 453–460.

Oswald, A., M. Streubel, U. Ljungberg, J. Hermans, K. Eskins, and P. Westhoff. 1990. “Differential Biogenesis of Photosystem-II in Mesophyll and Bundle-Sheath Cells of ‘Malic’ Enzyme NADP(+)-Type C4 Plants. A Comparative Protein and RNA Analysis.” European Journal of Biochemistry 190 (1): 185–194.

Rao, Xiaolan, and Richard A. Dixon. 2016. “The Differences between NAD-ME and NADP-ME Subtypes of C4 Photosynthesis: More than Decarboxylating Enzymes.” Frontiers in Plant Science 7 (October): 215899.

Retta, Moges A., Xinyou Yin, Quang Tri Ho, et al. 2023. “The Role of Chloroplast Movement in C4 Photosynthesis: A Theoretical Analysis Using a Three-Dimensional Reaction-Diffusion Model for Maize.” Journal of Experimental Botany 74 (14): 4125–4142.

Rosar, Christian, Kerstin Kanonenberg, Arun M. Nanda, et al. 2012. “The Leaf Reticulate Mutant dov1 Is Impaired in the First Step of Purine Metabolism.” Molecular Plant 5 (6): 1227–1241.

Sage, Rowan F. 2004. “The Evolution of C Photosynthesis.” The New Phytologist 161 (2): 341–370.

Sage, Rowan F., and Athena D. McKown. 2006. “Is C4 Photosynthesis Less Phenotypically Plastic than C3 Photosynthesis?” Journal of Experimental Botany 57 (2): 303–317.

Sage, Rowan F., Tammy L. Sage, and Ferit Kocacinar. 2012. “Photorespiration and the Evolution of C4 Photosynthesis.” Annual Review of Plant Biology 63 (January): 19–47.

Sage, Rowan F., Roxana Khoshravesh, and Tammy L. Sage. 2014. “From Proto-Kranz to C4 Kranz: Building the Bridge to C4 Photosynthesis.” Journal of Experimental Botany 65 (13): 3341–3356. 10.1093/jxb/eru180

Schlüter, Urte, Jacques W. Bouvier, Ricardo Guerreiro, et al. 2023. “*Brassicaceae* Display Variation in Efficiency of Photorespiratory Carbon-Recapturing Mechanisms.” Journal of Experimental Botany, July 1, erad250.

Schlüter, Urte, and Andreas P. M. Weber. 2020. “Regulation and Evolution of C Photosynthesis.” Annual Review of Plant Biology 71 (April): 183–215.

Schreier, Tina B., Karin H. Müller, Simona Eicke, Christine Faulkner, Samuel C. Zeeman, and Julian M. Hibberd. 2022. “Plasmodesmal Connectivity in C4 Gynandropsis Gynandra Is Induced by Light and Dependent on Photosynthesis.” bioRxiv. 10.1101/2022.12.07.519530.

Schreier, Tina B., Christian Paolo Balahadia, and Florence R. Danila. 2026. “Cell-to-Cell Connectivity: A Future Target for Crop Improvement.” Journal of Experimental Botany 77 (3): 697–713.

Schuler, Mara L., Otho Mantegazza, and Andreas P. M. Weber. 2016. “Engineering C4 Photosynthesis into C3 Chassis in the Synthetic Biology Age.” The Plant Journal 87 (1): 51–65.

Shameer, Sanu, Kambiz Baghalian, C. Y. Maurice Cheung, R. George Ratcliffe, and Lee J. Sweetlove. 2018. “Computational Analysis of the Productivity Potential of CAM.” Nature Plants 4 (3): 165–171.

Sheen, J. Y., R. T. Sayre, and L. Bogorad. 1987. “Differential Expression of Oxygen-Evolving Polypeptide Genes in Maize Leaf Cell Types.” Plant Molecular Biology 9 (3): 217–226.

Sundermann, Esther M., Martin J. Lercher, and David Heckmann. 2021. “Modeling Photosynthetic Resource Allocation Connects Physiology with Evolutionary Environments.” Scientific Reports 11 (1): 15979.

Takabayashi, Atsushi, Masahiro Kishine, Kozi Asada, Tsuyoshi Endo, and Fumihiko Sato. 2005. “Differential Use of Two Cyclic Electron Flows around Photosystem I for Driving CO2-Concentration Mechanism in C4 Photosynthesis.” Proceedings of the National Academy of Sciences 102 (46): 16898–16903.

Tobin, A. K., and T. Yamaya. 2001. “Cellular Compartmentation of Ammonium Assimilation in Rice and Barley.” Journal of Experimental Botany 52 (356): 591–604.

Töpfer, Nadine, Thomas Braam, Sanu Shameer, R. George Ratcliffe, and Lee J. Sweetlove. 2020. “Alternative Crassulacean Acid Metabolism Modes Provide Environment-Specific Water-Saving Benefits in a Leaf Metabolic Model.” The Plant Cell 32 (12): 3689–3705.

Weber, Andreas P. M., and Susanne von Caemmerer. 2010. “Plastid Transport and Metabolism of C3 and C4 Plants — Comparative Analysis and Possible Biotechnological Exploitation.” Current Opinion in Plant Biology 13 (3): 256–264.

Xiao, Yi, Jen Sloan, Chris Hepworth, et al. 2023. “Defining the Scope for Altering Rice Leaf Anatomy to Improve Photosynthesis: A Modelling Approach.” The New Phytologist 237 (2): 441–453.

Yin, Xinyou, and Paul C. Struik. 2021. “Exploiting Differences in the Energy Budget among C Subtypes to Improve Crop Productivity.” The New Phytologist 229 (5): 2400–2409.

Yin, Xinyou, Florian A. Busch, Paul C. Struik, and Thomas D. Sharkey. 2021. “Evolution of a Biochemical Model of Steady-State Photosynthesis.” Plant, Cell & Environment 44 (9): 2811–2837.

