## Supplementary Methods for "Metabolic modeling links leaf anatomy to environment-specific benefits on the C3-C4 spectrum"

[‡]both authors contributed equally

### Supplementary Methods

#### 1. Model Reconstruction

##### Core model of plant primary metabolism

We used a core model of plant metabolism (Shameer et al. 2018) with a representative plant biomass composition encompassing carbohydrates, cell wall constituents, lipids, and soluble metabolites from (de Oliveira Dal'Molin et al. 2010; Shameer et al. 2018). A two-cell mesophyll (M) -bundle sheath (B) system was constructed by duplicating the metabolic network for each cell type and connecting them via plasmodesmatal metabolite transport.

##### Naming conventions

In the model each reaction and metabolite is associated with a unique identifier following established naming conventions. Metabolite IDs are structured as {Metabolite Name}_{Compartment}. Exchange reactions between the model and extracellular space follow the convention {Reaction Name}_tx. Transport reactions between subcellular compartments use the structure {Metabolite Name}_{Origin Compartment}{Destination Compartment}. All other reactions follow the same convention as metabolites: {Reaction Name}_{Compartment}. Metabolite and reaction names are derived from MetaCyc (Caspi et al. 2020), while compartments are abbreviated using one or two letters (e.g., mitochondria - m; plastid - p; cytosol - c). In the two-cell-type model cell type prefixes were added to each metabolite and reaction ID resulting in the structure {Cell Type}_{ID}. Cell type prefixes are [B] for bundle sheath and [M] for mesophyll cells. Transport reactions of cytosolic metabolites between cell types were named using the exchange reaction convention with an added prefix: {[MB]}_{Metabolite Name}_tx.

##### Added reactions

Transport reactions absent from the base model were added to enable correct C4 flux distribution (Table S1). A malate/pyruvate shuttle was incorporated to facilitate pyruvate export from the BS plastid during the C4 cycle, and a pyruvate/proton symporter was added to enhance pyruvate transport to the plastid as in (Blätke and Bräutigam 2019).

**Table S1: Transport reactions added to the model to enable the C4 cycle.** Reaction formulas show metabolite exchange between cytosol (c) and plastid (p) compartments.

| Reaction Name | Reaction ID | Reaction Formula | Reference |
|---|---|---|---|
| Malate/Pyruvate transporter | PYR_MAL_pc | MAL_c + PYRUVATE_p ↔ MAL_p + PYRUVATE_c | (Linka and Weber 2010) |
| Proton-mediated Pyruvate symporter | PYR_H_pc | PROTON_c + PYRUVATE_c ↔ PROTON_p + PYRUVATE_p | (Caspi et al. 2020) |

#### Constraints on exchange reactions

The directionality of exchange reactions was set to provide biologically realistic flux solutions as summarised in Table S2. Uptake of $NH_4^+$ was blocked to prevent preferential uptake over $NO_3^-$. Gas exchange with the intracellular air space was restricted to the mesophyll and nutrient exchange with the vasculature was limited to bundle sheath cells as summarised in Table S3.

**Table S2: Upper and lower bounds of exchange reactions.**

| Reaction ID | Lower Bound | Upper Bound |
|---|---|---|
| H2O_tx | -1000 | 1000 |
| NH4_tx | 0 | 0 |
| Nitrate_tx | 0 | 1000 |
| Pi_tx | 0 | 1000 |
| SO4_tx | 0 | 1000 |
| O2_tx | -1000 | 1000 |
| Sucrose_tx | -1000 | 0 |
| GLC_tx | -1000 | 0 |
| Photon_tx | 0 | 1000 |
| unlProtHYPO_c | 0 | 0 |

**Table S3: Upper and lower bounds of exchange reactions specific to each cell-type.**

| Reaction ID | Lower Bound | Upper Bound |
|---|---|---|
| [B]_CO2_tx | 0 | 0 |
| [M]_CO2_tx | 0 | 20 |
| [M]_Nitrate_tx | 0 | 0 |
| [M]_SO4_tx | 0 | 0 |
| [M]_H2O_tx | 0 | 0 |
| [M]_Ca_tx | 0 | 0 |
| [M]_Mg_tx | 0 | 0 |
| [M]_Pi_tx | 0 | 0 |

#### Constraints on plasmodesmatal transport reactions

The two-cell type model allows for the exchange of 194 cytosolic metabolites between the M and BS cells, including amino acids, sugars, single phosphorylated sugar, carboxylic acids, glyceric acids, glycolate, glycerate, glyceraldehyde-3-phosphate, di-hydroxyacetone-phosphate, and $CO_2$. A further 47 cytosolic metabolites were excluded, based on the work of (Blätke and Bräutigam 2019) as shown in Table S4.

**Table S4: List of cytosolic metabolites blocked from exchange between the M and BS cells.**

| Metabolite Name | Metabolite ID | Metabolite Name | Metabolite ID |
|---|---|---|---|
| hydrogen sulfide ($H_2S$) | HS_c | oxaloacetate ($C_4H_2O_5$) | OXALACETIC_ACID_c |

| Metabolite Name | Metabolite ID | Metabolite Name | Metabolite ID |
|---|---|---|---|
| fructose 1,6-bisphosphate ($C_6H_{14}O_{12}P_2$) | FRUCTOSE_16_DIPHOSPHATE_c | hydrogencarbonate ($HCO_3$) | HCO3_c |
| 3-phospho-D-glyceroyl phosphate ($C_3H_4O_{10}P_2$) | DPG_c | UTP ($C_9H_{11}N_2O_{15}P_3$) | UTP_c |
| proton ($H^+$) | PROTON_c | UDP ($C_9H_{11}N_2O_{12}P_2$) | UDP_c |
| acetaldehyde ($C_2H_4O$) | ACETALD_c | UDP-α-D-glucose ($C_{15}H_{22}N_2O_{17}P_2$) | UDP_GLUCOSE_c |
| acetate ($C_2H_3O_2$) | ACET_c | ATP ($C_{10}H_{12}N_5O_{13}P_3$) | ATP_c |
| 5,10-methenyl tetrahydrofolate ($C_{20}H_{22}N_7O_6$) | 5_10_METHENYL_THF_c | ADP ($C_{10}H_{12}N_5O_{10}P_2$) | ADP_c |
| 5-methyltetrahydrofolate ($C_{20}H_{25}N_7O_6$) | 5_METHYL_THF_c | AMP ($C_{10}H_{12}N_5O_7P$) | AMP_c |
| L-homocysteine ($C_4H_9NO_2S$) | HOMO_CYS_c | IMP ($C_{10}H_{11}N_4O_8P$) | IMP_c |
| S-adenosyl-L-homocysteine ($C_{14}H_{20}N_6O_5S$) | ADENOSYL_HOMO_CYS_c | XMP ($C_{10}H_{11}N_4O_9P$) | XANTHOSINE_5_PHOSPHATE_c |
| O-acetyl-L-serine ($C_5H_9NO_4$) | ACETYLSERINE_c | GTP ($C_{10}H_{12}N_5O_{14}P_3$) | GTP_c |
| tetrahydrofolate ($C_{19}H_{23}N_7O_6$) | THF_c | GDP ($C_{10}H_{12}N_5O_{11}P_2$) | GDP_c |
| adenosine ($C_{10}H_{13}N_5O_4$) | ADENOSINE_c | GMP ($C_{10}H_{12}N_5O_8P$) | GMP_c |
| maltose ($C_{12}H_{22}O_{11}$) | MALTOSE_c | CDP ($C_9H_{12}N_3O_{11}P_2$) | CDP_c |
| coenzyme A ($C_{21}H_{32}N_7O_{16}P_3S$) | CO_A_c | dUMP ($C_9H_{11}N_2O_8P$) | DUMP_c |
| γ-L-glutamyl 5-phosphate ($C_5H_8NO_7P$) | L_GLUTAMATE_5_P_c | dTMP ($C_{10}H_{13}N_2O_8P$) | dTMP_c |
| acetyl-CoA ($C_{23}H_{34}N_7O_{17}P_3S$) | ACETYL_COA_c | dTDP ($C_{10}H_{13}N_2O_{11}P_2$) | DTDP_c |
| cellulose ($(C_6H_{10}O_5)$)□ | CELLULOSE_c | GTP ($C_{10}H_{12}N_5O_{14}P_3$) | GTP_c |
| L-glutamate-5-semialdehyde ($C_5H_9NO_3$) | L_GLUTAMATE_GAMMA_SEMIALDEHYDE_c | dTTP ($C_{10}H_{13}N_2O_{14}P_3$) | DTTP_c |
| S-adenosyl-L-methionine ($C_{15}H_{23}N_6O_5S$) | S_ADENOSYLMETHIONINE_c | NAD⁺ ($C_{21}H_{26}N_7O_{14}P_2$) | NAD_c |
| diphosphate ($H_4O_7P_2$) | PPI_c | NADH ($C_{21}H_{27}N_7O_{14}P_2$) | NADH_c |
| (S)-1-pyrroline-5-carboxylate ($C_5H_6NO_2$) | L_DELTA1_PYRROLINE_5_CARBOXYLATE_c | NADP⁺ ($C_{21}H_{25}N_7O_{17}P_3$) | NADP_c |
| ammonium ($NH_4^+$) | AMMONIUM_c | NADPH ($C_{21}H_{26}N_7O_{17}P_3$) | NADPH_c |
| carbon dioxide ($CO_2$) | CARBON_DIOXIDE_c | | |

#### Constraints on internal reactions

Plastoquinol oxidase, nucleotide translocator (NTT), and uncoupled pyruvate transport were blocked as in (Blätke and Bräutigam 2019). The ATP maintenance cost reaction was constrained to only allow for ATP consumption. In plant photosynthetic tissues, the Ferredoxin: NADP+ reductase (FNR) enzyme primarily works in the direction to transfer electrons from ferredoxin to NADPH (Mulo 2011) and the reaction was constrained accordingly. Pyruvate orthophosphate dikinase (PPDK) occurs mostly in M cell plastids (Sage 2004). All added internal constraints are detailed in Table S5.

**Table S5: List of constraints for internal reactions.**

| Reaction ID | Lower bound | Upper bound | Reference |
|---|---|---|---|
| **Both cell types** | | | |
| Plastoquinol_Oxidase_p | 0 | 0 | (Josse et al. 2000) |
| ATP_ADP_Pi_pc | 0 | 0 | (Reiser et al. 2004) |
| PYRUVATE_pc | 0 | 0 | (Blätke and Bräutigam 2019) |
| ATPase_tx | 0 | 1000 | - |
| 1_PERIOD_18_PERIOD_1_PERIOD_2_RXN_p | 0 | 1000 | (Mulo 2011) |
| **Mesophyll cell** | | | |
| [M]_PYRUVATEORTHOPHOSPHATE_DIKINASE_RXN_c | 0 | 0 | (Sage 2004) |

Initial FBA solutions showed no $C_4$ cycle activity. Traceback of C metabolism revealed that the $C_4$ cycle was bypassed through the reverse usage of TCA cycle and urea degradation reactions. This was corrected by adjusting the directionality of these reactions to their physiological context (Table S6).

**Table S6: List of constraints to alternative decarboxylation pathways.**

| Reaction ID | Lower bound | Upper bound |
|---|---|---|
| [B]_UREASE_RXN_c | 0 | 0 |
| [M]_UREASE_RXN_c | 0 | 1000 |
| [B]_CARBAMATE_KINASE_RXN_p | 0 | 1000 |
| [M]_CARBAMATE_KINASE_RXN_p | 0 | 1000 |
| [B]_ISOCITDEH_RXN_m | 0 | 1000 |
| [M]_ISOCITDEH_RXN_m | 0 | 1000 |
| [B]_ISOCITDEH_RXN_c | 0 | 1000 |
| [M]_ISOCITDEH_RXN_c | 0 | 1000 |
| [B]_ISOCITRATE_DEHYDROGENASE_NAD_RXN_m | 0 | 1000 |
| [M]_ISOCITRATE_DEHYDROGENASE_NAD_RXN_m | 0 | 1000 |

#### 2. Anatomical constraints

##### Anatomical data collection

Light or electron microscopy images of leaf cross-sections (200 samples) and paradermal sections (2 samples) from 84 mono- and dicot species (25 C3, 28 C3-C4 intermediates, 31 C4) were extracted from 18 publications, resulting in a total of 202 sample images (Notebook Anatomical_Data_Analysis). For each sample, M and BS cell type-enriched volumes were quantified by measuring the widths of BS and M enriched volumes parallel to the leaf surface as illustrated in Figure S1. All measurements were performed using Fiji (ImageJ) version 1.54g.

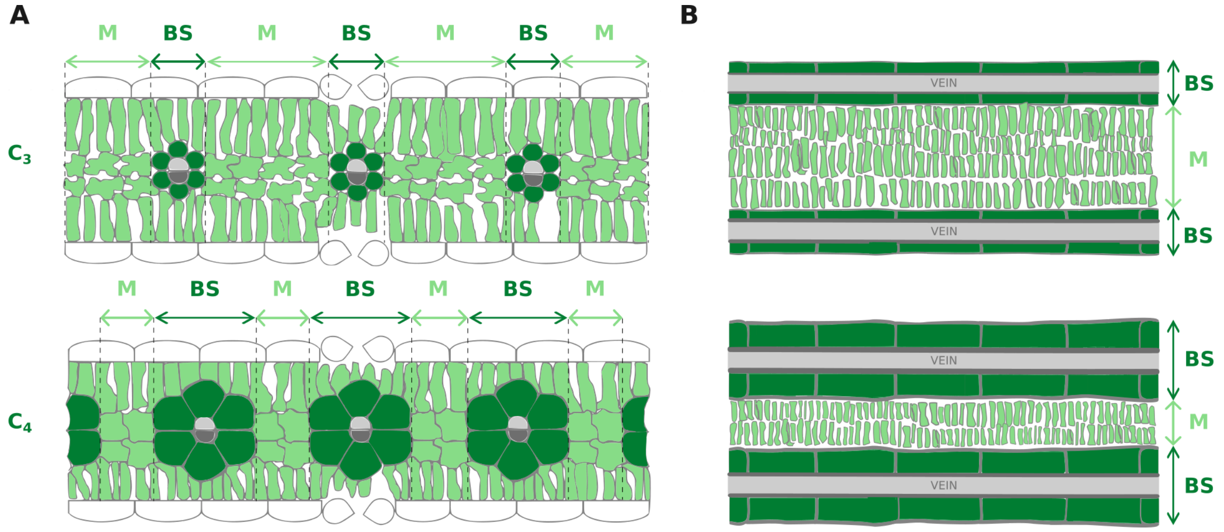


**Figure S1: Anatomical measurements of mesophyll and bundle sheath cell type-enriched volumes in leaf sections.** Bundle sheath cell-enriched volume width was defined as the diameter of the vascular bundle; mesophyll cell-enriched volume width was defined as the interveinal distance. All measurements were made parallel to the leaf epidermis. (A) Cross-sectional measurements; (B) Paradermal section measurements.

For each leaf section, individual cell type-enriched volume measurements were averaged to obtain sample-level values of bundle sheath ($W_{BS}$) and mesophyll ($W_M$) widths. Volume ratios were then calculated using the equation:

$$V_{\frac{BS}{M}} = \frac{L_{BS} \cdot W_{BS} \cdot D_{BS}}{L_M \cdot W_M \cdot D_M} = \frac{A_{BS}}{A_M} = \frac{W_{BS}}{W_M}$$

, where $L_{BS}$ and $L_M$ is the cell-type enriched volume's length and $D_{BS}$ and $D_M$ the thickness for M and B, which are both assumed to be equal for BS and M.

Outlier species within each photosynthetic type were identified and removed using Tukey's method with a 1.5× IQR multiplier. This resulted in interquartile ranges of 0.5 - 1.1 for C3, 0.7 - 1.7 for intermediates, and 1.2 - 2.2 for C4 species. The final species subset was visualised as a phylogenetic tree using the ETE3 toolkit in Python. The phylogengy was constructed using the NCBI Taxonomy database (Schoch et al. 2020). Branches were colored according to their photosynthetic type and $V_{BS/M}$ values represented as colour-coded ring segments (Figure S2).

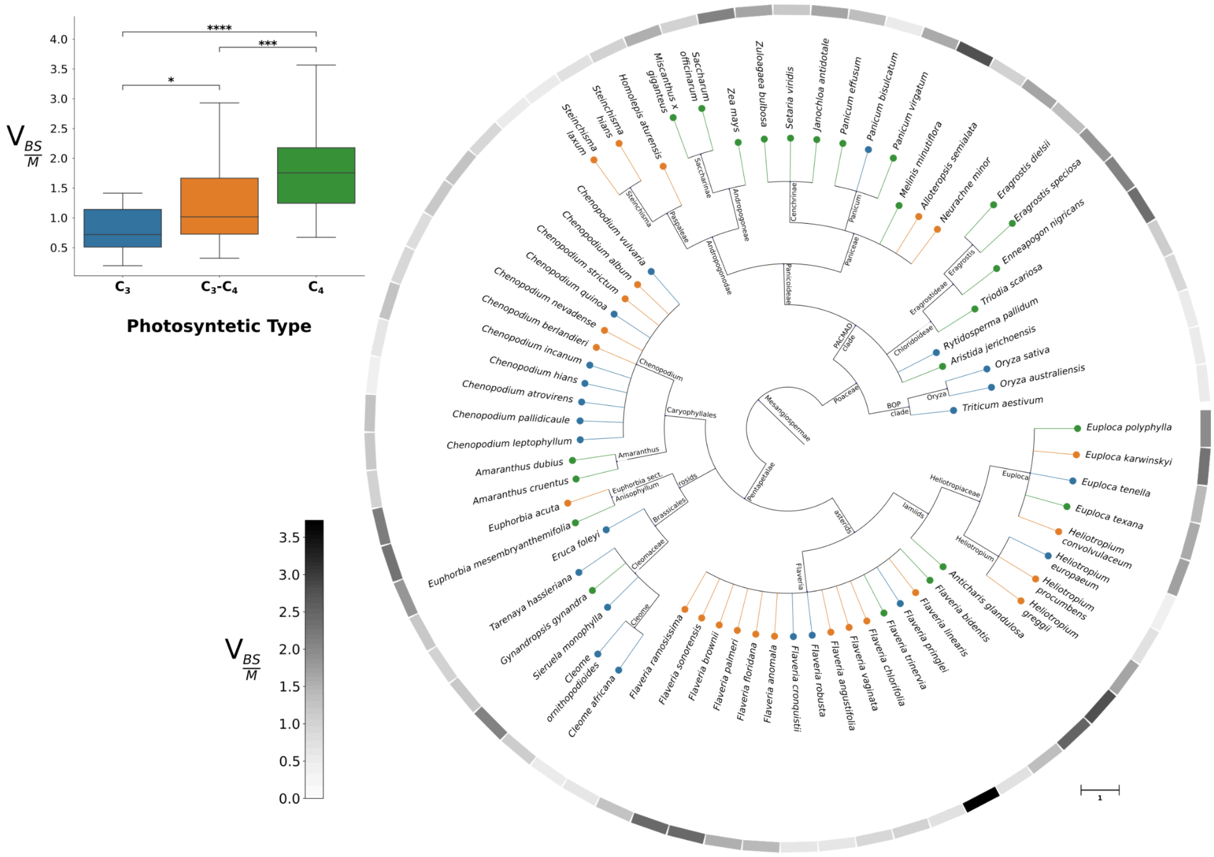


**Figure S2: Phylogenetic tree and summary of mesophyll and bundle sheath volume ratio data collection.** Leaf colour represents the photosynthetic type of the given species. $C_3$ species are represented in blue, $C_3$-$C_4$ intermediates are represented in orange, and $C_4$ species are represented in green. The ratio of the bundle sheath and mesophyll cell-enriched volumes is represented as colour-coded ring segments. Volume ratios for the $C_3$, $C_3$-$C_4$ and $C_4$ photosynthetic types are shown in the boxplot. P-values were obtained through the Mann–Whitney U test, where one * represents a p-value between 0.05 and 0.01, ** between 0.01 and 0.001 and *** between 0.001 and 0.0001.

#### Scaling fluxes according to the cell-type enriched volumes

We consider the M and BS as interacting subsystems and thus model their fluxes per cell type-enriched volume and not per leaf volume. As a consequence, exchange fluxes from the environment and between the cell types need to be scaled according to the cell type volumes. Note, that we assume that the volume ratio of BS:M is equal to the area ratio of BS:M, as we assume the leaf thickness to be equal for both volume approximations. This is important as fluxes are given per $m^2$. For a given $V_{BS/M}$ we can then calculate $V_M$ and $V_{BS}$ using the following equations:

$$V_{BS} = \frac{V_{\frac{BS}{M}}}{\left(1 + V_{\frac{BS}{M}}\right)}$$

$$V_{M} = \frac{1}{\left(1 + V_{\frac{BS}{M}}\right)}$$

We use these values to scale exchange fluxes with the environment and between the cell types as shown in Figure S3. For example, for nitrate uptake into the BS cells from the vasculature, subsequent transport to the M and (theoretical) exchange with the intracellular air space from the M, the fluxes would be normalized using the following equations.

$$[B]\_Nitrate\_tx: \quad 1[B]\_Nitrate\_e \leftrightarrow \frac{1}{V_{BS}} [B]\_Nitrate\_c$$

$$[M]\_Nitrate\_tx: \quad 1[M]\_Nitrate\_e \leftrightarrow \frac{1}{V_{M}} [M]\_Nitrate\_c$$

$$[MB]\_Nitrate\_tx: \quad \frac{1}{V_{M}} [M]\_Nitrate\_c \leftrightarrow \frac{1}{V_{BS}} [B]\_Nitrate\_c$$

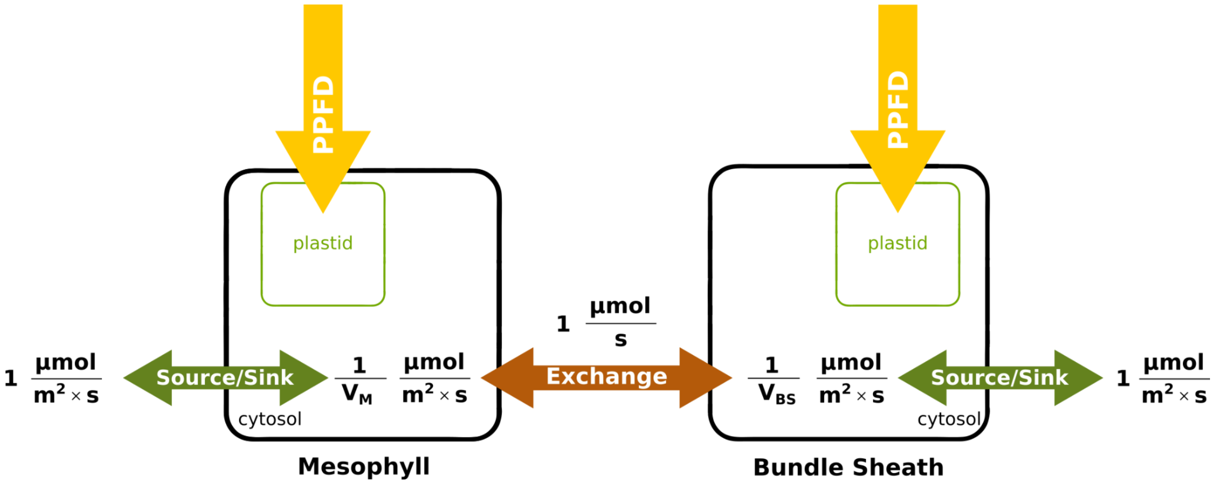


**Figure S3: Scaling exchange reaction fluxes according to M and BS cell enriched leaf volumes.** Flux entering each cell-type submodel is normalised by dividing it by the value of the respective cell-type enriched volume. When flux is exchanged between the cell types, it is first divided by the volume of the originating cell type and then divided by the volume of the destination cell type.

To illustrate this, consider 1m$^2$ of leaf supplied with 1 µmol nitrate s$^{-1}$ from the vasculature. Let us also assume a BS:M ratio of 1:3, i.e. 25% of the leaf area is enriched in BS cells and 75% in M cells. Import of this nitrate into the BS corresponds to an uptake rate of 4 µmol m$^{-2}$ s$^{-1}$ per BS cell-type enriched area (1 µmol s$^{-1}$ / 0.25 m$^2$).

If this nitrate is then transported to the M, the exchange flux from the BS is likewise 4 µmol m$^{-2}$ s$^{-1}$ per BS-enriched area. Upon entering the M, this translates into an uptake flux of 1.33 µmol m$^{-2}$ s$^{-1}$ per M-enriched area (1 µmol s$^{-1}$ / 0.75 m$^2$). In the purely theoretical case that this flux was released into the intercellular air space, it would amount to 1 µmol m$^{-2}$ leaf s$^{-1}$ (1.33 µmol m$^{-2}$ s$^{-1}$ M x 0.75 m$^2$ M).

Note that light uptake does not follow that logic and light uptake is modelled independently for each cell type. For instance, for a leaf area of 1m$^2$ a light intensity of 1000 µmol photons m$^{-2}$ s$^{-1}$ and a $V_{BS/M}$ of 1:3, this means that a maximum of 750 µmol photons s$^{-1}$ can enter the M cell-enriched area and a maximum of 250 µmol photons s$^{-1}$ can enter the BS cell-enriched area because this makes in total 1000 µmol photons m$^{-2}$ s$^{-1}$.

### Supplementary Results

##### Predicted photosynthetic types without plasmodesdamtal flux weighting

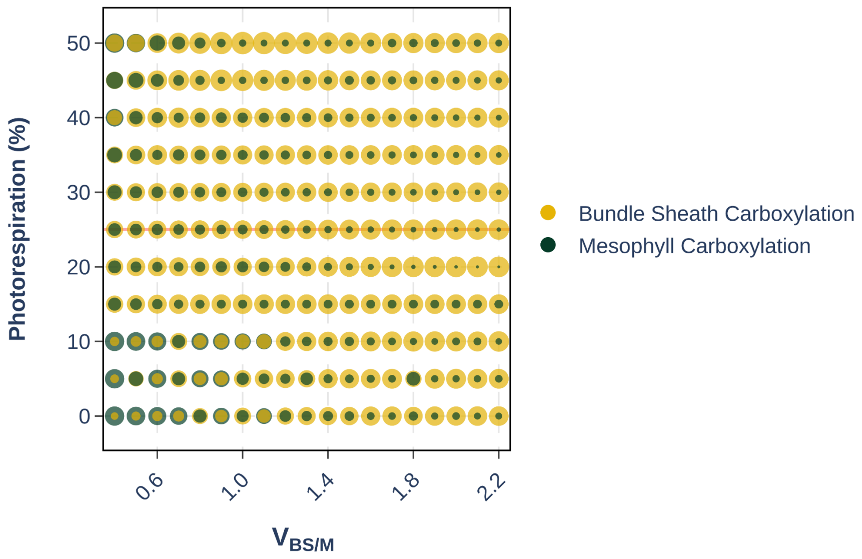


**Figure S4: Predicted photosynthetic types for different levels of photorespiration and bundle sheath to mesophyll volume ($V_{BS/M}$) ratios without plasmodesmatal flux weighting.** Each bubble represents an individual flux solution. Bubble colours indicate rubisco activity in the mesophyll or bundle sheath and bubble size encodes rubisco flux value. Overlapping bubbles indicate concurrent activity of both reactions. The red line marks the rubisco $v_c/v_O$ ratio of 3:1 assumed under ambient conditions.

**Table S7: Thermodynamic estimates for non-canonical $CO_2$-releasing reactions in pre-C2 flux modes.** Listed are equilibrium-standard transformed Gibbs energy (ΔrG′°) values obtained from AraCyc (Mueller et al. 2003), ΔrG′° and transformed Gibbs energy (ΔrG′) values obtained using Equilibrator (Beber et al. 2022) for comparison as well as a qualitative feasibility call.

| Model reaction ID | Reaction | Enzyme | Compartment (pH) | AraCyc ΔrG′° (kcal ·mol⁻¹)/(kJ ·mol⁻¹) | Equilibrator ΔrG′° (kJ·mol⁻¹, mean ± SD) | Feasibility |
|---|---|---|---|---|---|---|
| **Amino acid biosynthesis** | | | | | | |
| [B]_DIAMINOPIMDECARB_RXN_p | meso-2,6-diaminopimelate → L-lysine + $CO_2$ | Diaminopimelate decarboxylase (EC 4.1.1.20) | Plastid (8.0) | −3.09 / −12.93 | −12.52 ± 3.73 | Favorable |
| [B]_ACETOOHBUTSYN_RXN_p | 2-oxobutanoate + pyruvate + $H^+$ → 2-aceto-2-hydroxybutyrate + $CO_2$ | Acetolactate synthase (EC 2.2.1.6) | Plastid (8.0) | −9.70 / −40.57 | −15.73 ± 4.91 | Favorable |
| [B]_RXN_5682_p | L-arogenate + $NADP^+$ → L-tyrosine + $CO_2$ + NADPH | Arogenate dehydrogenase (EC 1.3.1.78) | Plastid (8.0) | −17.49 / −73.18 | −20.28 ± 11.06 | Favorable |
| [B]_RXN_7800_p | CPD-7100 + $H^+$ → 2-keto-4-methylpentanoate + $CO_2$ | β-isopropylmalate dehydrogenase (EC 1.1.1.85) | Plastid (8.0) | −1.84 / −7.70 | −11.30 ± 4.33 | Favorable |
| [B]_ACETOLACTSYN_RXN_p | 2 pyruvate → 2-acetolactate + $CO_2$ | Acetolactate synthase (EC 2.2.1.6) | Plastid (8.0) | −9.49 / −39.71 | −24.63 ± 3.73 | Favorable |

| Model reaction ID | Reaction | Enzyme | Compartment (pH) | AraCyc ΔrG'° (kcal ·mol$^{-1}$)/(kJ ·mol$^{-1}$) | Equilibrator ΔrG'° (kJ·mol$^{-1}$, mean ± SD) | Feasibility |
|---|---|---|---|---|---|---|
| [B]_CARBOXYCYCLOHEXADIENYL_DEHYDRATASE_RXN_p | L-arogenate → L-phenylalanine + $CO_2$ + $H_2O$ | Arogenate dehydratase (EC 4.2.1.91) | Plastid (8.0) | −26.89 / −112.51 | −71.39 ± 11.07 | Favorable |
| **Fatty acid biosynthesis** | | | | | | |
| [B]_2.3.1.180_RXN_p | acetyl-CoA + malonyl-ACP + $H^+$ → acetoacetyl-ACP + $CO_2$ + CoA | KAS III / FabH (EC 2.3.1.180) | Plastid (8.0) | −27.41 / −114.68 | n/a | Favorable |
| [B]_RXN_9648_p | butanoyl-CoA + malonyl-CoA → 3-oxo-hexanoyl-CoA + $CO_2$ | β-ketoacyl-ACP synthase | Plastid (8.0) | −15.26 / −63.85 | n/a | Favorable |
| [B]_RXN_9650_p | hexanoyl-CoA + malonyl-CoA → 3-oxo-octanoyl-CoA + $CO_2$ | β-ketoacyl-ACP synthase | Plastid (8.0) | −15.26 / −63.85 | n/a | Favorable |
| [B]_RXN_9651_p | octanoyl-CoA + malonyl-CoA → 3-oxo-decanoyl-CoA + $CO_2$ | β-ketoacyl-ACP synthase | Plastid (8.0) | −15.26 / −63.85 | n/a | Favorable |
| [B]_RXN_9652_p | decanoyl-CoA + malonyl-CoA → 3-oxo-dodecanoyl-CoA + $CO_2$ | β-ketoacyl-ACP synthase | Plastid (8.0) | −15.26 / −63.85 | n/a | Favorable |
| [B]_RXN_9653_p | dodecanoyl-CoA + malonyl-CoA → 3-oxo-myristoyl-CoA + $CO_2$ | β-ketoacyl-ACP synthase | Plastid (8.0) | −15.26 / −63.85 | n/a | Favorable |
| [B]_RXN_9654_p | myristoyl-CoA + malonyl-CoA → 3-oxo-palmitoyl-CoA + $CO_2$ | β-ketoacyl-ACP synthase | Plastid (8.0) | −15.26 / −63.85 | n/a | Favorable |
| **Central carbon metabolism** | | | | | | |
| [B]_PYRUVDEH_RXN_p | pyruvate + CoA + $NAD^+$ → acetyl-CoA + $CO_2$ + NADH | Pyruvate dehydrogenase (plastid) | Plastid (8.0) | −9.98 / −41.76 | −34.04 ± 3.22 | Favorable |
| [B]_PYRUVDEH_RXN_m | pyruvate + CoA + $NAD^+$ → acetyl-CoA + $CO_2$ + NADH | Pyruvate dehydrogenase (mitochondrial) | Mitochondrion (7.8) | −9.98 / −41.76 | −34.34 ± 3.22 | Favorable |
| [B]_ISOCITDEH_RXN_c | isocitrate + $NADP^+$ → 2-oxoglutarate + $CO_2$ + NADPH | Isocitrate dehydrogenase (EC 1.1.1.42) | Cytosol (7.2) | 0.38 / 1.57 | 4.64 ± 3.81 | Favorable |

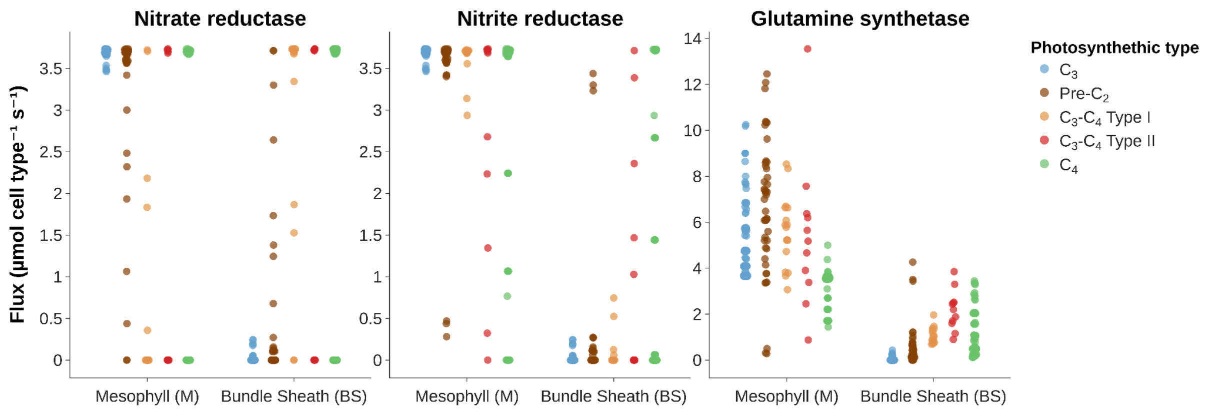

**Figure S5: Cell-type-specific flux distributions for key nitrogen assimilation enzymes across photosynthetic types.** Mesophyll (M) and bundle sheath (B) fluxes are shown for nitrate reductase, nitrite reductase, and total glutamine synthetase. Each point represents an individual flux balance solution obtained from a parameter scan of BS-M anatomies between 0.1 - 2.2 and photorespiratory levels (5–50% of total rubisco flux is oxygenation reaction flux).

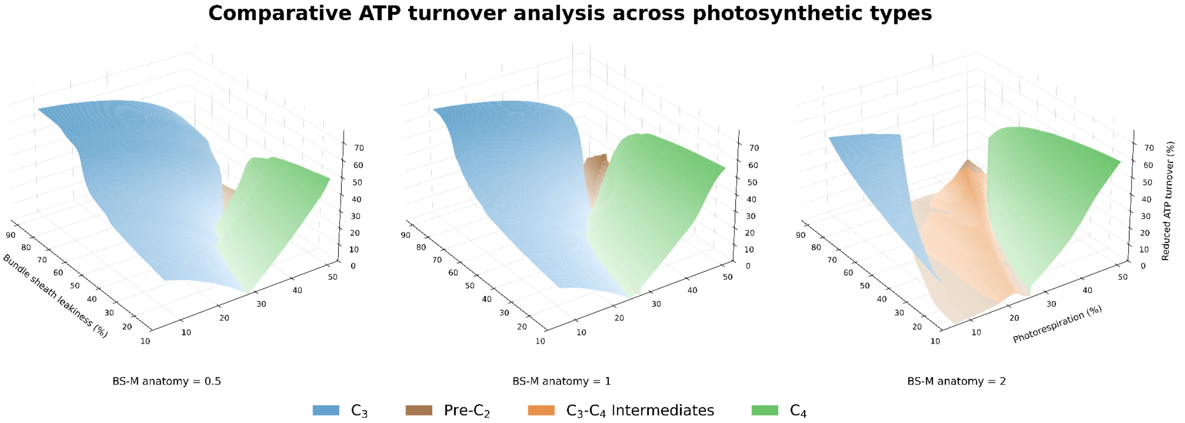


**Figure S6: Photosynthetic optimality in dependence of bundle sheath leakage and photorespiration for different BS-M anatomies.** Metabolic flux patterns were classified as C3, pre-C2, C3-C4 intermediate, or C4 photosynthesis. For flux patterns classified as C3-C4 intermediates, energetic efficiency, as measured by ATP turnover, was benchmarked against two reference scenarios: a C3 scenario confining rubisco activity to the mesophyll, and a C4 scenario confining rubisco activity to the bundle sheath. The reduction in ATP turnover relative to the second most efficient reference scenario was then calculated and plotted. For flux patterns classified as C3 or C4, the reduction relative to the other reference scenario was plotted.

#### Bibliography


Beber, Moritz E., Mattia G. Gollub, Dana Mozaffari, et al. 2022. “eQuilibrator 3.0: A Database Solution for Thermodynamic Constant Estimation.” *Nucleic Acids Research* 50 (D1): D603–D609.

Blätke, Mary-Ann, and Andrea Bräutigam. 2019. “Evolution of C4 Photosynthesis Predicted by Constraint-Based Modelling.” *eLife* 8 (December). https://doi.org/10.7554/eLife.49305.

Caspi, Ron, Richard Billington, Ingrid M. Keseler, et al. 2020. “The MetaCyc Database of Metabolic Pathways and Enzymes - a 2019 Update.” *Nucleic Acids Research* 48 (D1): D445–D453.

Josse, E. M., A. J. Simkin, J. Gaffé, A. M. Labouré, M. Kuntz, and P. Carol. 2000. “A Plastid Terminal Oxidase Associated with Carotenoid Desaturation during Chromoplast Differentiation.” *Plant Physiology* 123 (4): 1427–1436.

Linka, Nicole, and Andreas P. M. Weber. 2010. “Intracellular Metabolite Transporters in Plants.” *Molecular Plant* 3 (1): 21–53.

Mueller, Lukas A., Peifen Zhang, and Seung Y. Rhee. 2003. “AraCyc: A Biochemical Pathway Database for Arabidopsis.” *Plant Physiology* 132 (2): 453–460.

Mulo, Paula. 2011. “Chloroplast-Targeted Ferredoxin-NADP(+) Oxidoreductase (FNR): Structure, Function and Location.” *Biochimica et Biophysica Acta* 1807 (8): 927–934.

Oliveira Dal’Molin, Cristiana Gomes de, Lake-Ee Quek, Robin William Palfreyman, Stevens Michael Brumbley, and Lars Keld Nielsen. 2010. “AraGEM, a Genome-Scale Reconstruction of the Primary Metabolic Network in Arabidopsis.” *Plant Physiology* 152 (2): 579–589.

Reiser, Jens, Nicole Linka, Lilia Lemke, Wolfgang Jeblick, and H. Ekkehard Neuhaus. 2004. “Molecular Physiological Analysis of the Two Plastidic ATP/ADP Transporters from Arabidopsis.” *Plant Physiology* 136 (3): 3524–3536.

Sage, Rowan F. 2004. “The Evolution of C4 Photosynthesis.” *The New Phytologist* 161 (2): 341–370.

Schoch, Conrad L., Stacy Ciufo, Mikhail Domrachev, et al. 2020. “NCBI Taxonomy: A Comprehensive Update on

Curation, Resources and Tools." *Database : The Journal of Biological Databases and Curation* 2020 (January). https://doi.org/10.1093/database/baaa062.

Shameer, Sanu, Kambiz Baghalian, C. Y. Maurice Cheung, R. George Ratcliffe, and Lee J. Sweetlove. 2018. "Computational Analysis of the Productivity Potential of CAM." *Nature Plants* 4 (3): 165–171.
